# Deep Mutational Engineering of broadly-neutralizing and picomolar affinity nanobodies to accommodate SARS-CoV-1 & 2 antigenic polymorphism

**DOI:** 10.1101/2021.12.07.471597

**Authors:** Adrien Laroche, Maria Lucia Orsini Delgado, Philippe Cuniasse, Steven Dubois, Raphaël Sierocki, Fabrice Gallais, Stéphanie Debroas, Laurent Bellanger, Stéphanie Simon, Bernard Maillère, Hervé Nozach

## Abstract

We report in this study the molecular engineering of nanobodies that bind with picomolar affinity to both SARS-CoV-1 and SARS-CoV-2 Receptor Binding Domains (RBD) and are highly neutralizing. We applied Deep Mutational Engineering to VHH72, a nanobody initially specific for SARS-CoV-1 RBD with little cross-reactivity to SARS-CoV-2 antigen. We first identified all the individual VHH substitutions that increase binding to SARS-CoV-2 RBD and then screened highly focused combinatorial libraries to isolate engineered nanobodies with improved properties. The corresponding VHH-Fc molecules show high affinities for SARS-CoV-2 antigens from various emerging variants and SARS-CoV-1, block the interaction between ACE2 and RBD and neutralize the virus with high efficiency. Its rare specificity across sarbecovirus relies on its peculiar epitope outside the immunodominant regions. The engineered nanobodies share a common motif of three amino acids, which contribute to the broad specificity of recognition. These nanobodies appears as promising therapeutic candidates to fight SARS-CoV-2 infection.

## Introduction

The severe acute respiratory syndrome coronavirus 2 (SARS-CoV-2) is the cause of the current COVID-19 pandemic, with devastating consequences in many countries. To date, vaccination of populations appears to be the most effective approach to control the magnitude of the epidemic and to effectively protect the health of individuals. In high-risk infected patients, passive immunization by infusion of monoclonal antibodies constitutes a very interesting and complementary approach to vaccination. Indeed, several studies demonstrated that early administration of monoclonal antibodies blocks entry of the virus in human cells by targeting the Spike protein and prevents progression to severe forms in patients ^1-3^.

Since early 2020, numerous studies have been dedicated to the development of monoclonal antibodies targeting the Spike protein and more particularly its Receptor Binding Domain (RBD) ^4^. Multiple SARS-CoV-2 antibodies described in the literature bind the receptor-binding motif (RBM) of the S protein i.e. the interaction site between the RBD domain and ACE2. These monoclonal antibodies often exhibit excellent affinities, very good ability to neutralize the SARS-CoV-2 virus, and a protective effect *in vivo*. Yet, most of these antibodies have a relatively narrow recognition spectrum, with a lack of recognition of RBD domains from other members of the sarbecovirus subgenus, including SARS-CoV-1 ^5^. Over the past year, new strains have emerged with mutations in the RBM affecting the transmissibility of the virus but also contributing to the escape from host immunity ^6-9^. Expansion of new viral variants in the overall population seems to be favoured by host immune pressure from prior infection ^10,11^ or to vaccination. Indeed, RBM is targeted by a large proportion of neutralizing antibodies and hence appears to be an immunodominant region. Many RBM residues are permissive to mutations, with a preserved binding to ACE2 but might affect the recognition by antibodies from patients ^12,13^ but also by therapeutic monoclonal antibodies and consequently alter their protective efficacy. Antibodies such as bamlanivimab or etesevimab, are subject to significant loss of affinity for emerging SARS-CoV-2 variants^12,14^ affecting their neutralization potency by a factor up to 1000 ^12,15^. Most of the antibodies have been produced by animal immunisation or B cell cloning from infected patients and hence are issued from the immunodominant B-cell repertoires, which are also involved in the selective pressure on the virus and thus participate to viral escape. This limitation illustrates the need of alternative strategies to develop antibodies with high neutralizing activity, broad spectrum of recognition of circulating sarbecoviruses (BnAbs) and limited sensitivity to immune escape adaptations.

Early after the beginning of the current pandemic, the proximity of the Spike protein sequences of the SARS-CoV-1 and SARS-CoV-2 strains led several teams to search for cross-reactive antibodies, from SARS-CoV-1 convalescent patients or by screening monoclonal antibody libraries obtained following the 2003 epidemic ^16-18^. Unfortunately, most monoclonal antibodies have restricted specificity for either SARS-CoV-1 or SARS-CoV-2 ^19^ and the ability to cross-neutralize both strains is a relatively uncommon feature ^20,21^. Sera of patients infected with either SARS-CoV-1 or SARS-CoV-2 show a poor cross-reactivity suggesting that common epitopes are rare. Rare examples of cross- reactive and neutralizing antibodies described in the literature include antibodies S309 ^17^ (from which sotrovimab is derived), antibodies S2H97^5^, ADG2^22^ or the VHH-Fc antibody rimteravimab, derived from the VHH72 nanobody ^18^.

VHH72 is a molecule originally resulting from a llama immunization with the SARS-CoV-1 Spike protein and which is able to recognize the corresponding SARS-CoV-2 antigen^18^. Binding affinity of VHH-72 to the SARS-CoV-2 RBD is moderate (∼39-60⍰nM), markedly weaker than that of the SARS- CoV-1 RBD (1.2 nM) ^18,23^. Despite relatively fast dissociation rates, this VHH is nonetheless able to neutralize SARS-CoV-2 pseudovirus when expressed as dimer or VHH-Fc.^18^ Published structure reveals that its epitope is relatively conserved within sarbecoviruses, and is located in the RBD core, distant from the ACE2 binding site, probably because of its involvement in the dynamics between “up” and “down” conformations of the spike protein ^23^. This VHH72 epitope meets therefore advantages of conservation across sarbecoviruses and limited permissivity to mutations but only exhibit a moderate affinity, which potentially impede its full potential for therapeutic applications.

We therefore engineered, in the present study, the VHH72 antibody by combining Deep Mutational Scanning and Yeast Surface Display to generate antibody candidates with greatly improved affinity and broad cross-reactivity for SARS-CoV-1 and SARS-CoV-2 antigens, and cross-neutralization of SARS-CoV-2 variants Alpha, Beta, Gamma and Delta.

## Results

For affinity maturation of the VHH-72 nanobody two successive mutational approaches were combined. We first used Deep Mutational Scanning to map permissive positions and identify substitutions demonstrating beneficial effects on antigen binding. We then generated focused combinatorial libraries with relatively modest size (<10^7^ clones) in which diversity is essentially restricted to beneficial or neutral substitutions. These libraries were then screened to identify clones combining different substitutions conferring them a substantial gain in affinity.

Two libraries encompassing all single amino changes in the VHH72 nanobody were designed with a single NNK degenerate codon for a given cDNA molecule, spanning respectively amino acids positions 1-59 and 60-125 (Fig 1A). A Yeast-Surface Display (YSD) system based on the Aga2p/Aga1p anchoring proteins allowed both protein expression and functional discrimination of clones based on their affinity for the SARS-CoV-2 RBD antigen (Fig 1B). The two libraries were cloned in the plasmid pNT VHH72, transformed in *S. cerevisiae* EBY100 and their expression was induced. To finely compare the RBD binding of clones harbouring mutations with VHH72, we introduced a small percentage (typically 5-10%) of cells expressing the parental clone along with intracellular egfp, using another plasmid (pSWG VHH72 efgp). In that manner, parental clones (in black) can be readily identified during the FACS analysis to precisely define adequate sorting gates (Fig 1C). Upon induction and incubation with biotinylated RBD SARS-CoV-2 antigen, cells displaying variants with strong surface expression levels and augmented RBD binding relatively to the parental clone were sorted (in red, Fig 1C). Overall, these sorted populations correspond to a relatively low percentage of cells expressing the VHH construct (respectively 5.5% and 11% in libraries L1 and L2), most mutants having similar or impaired RBD binding levels.

**Fig. 1.**
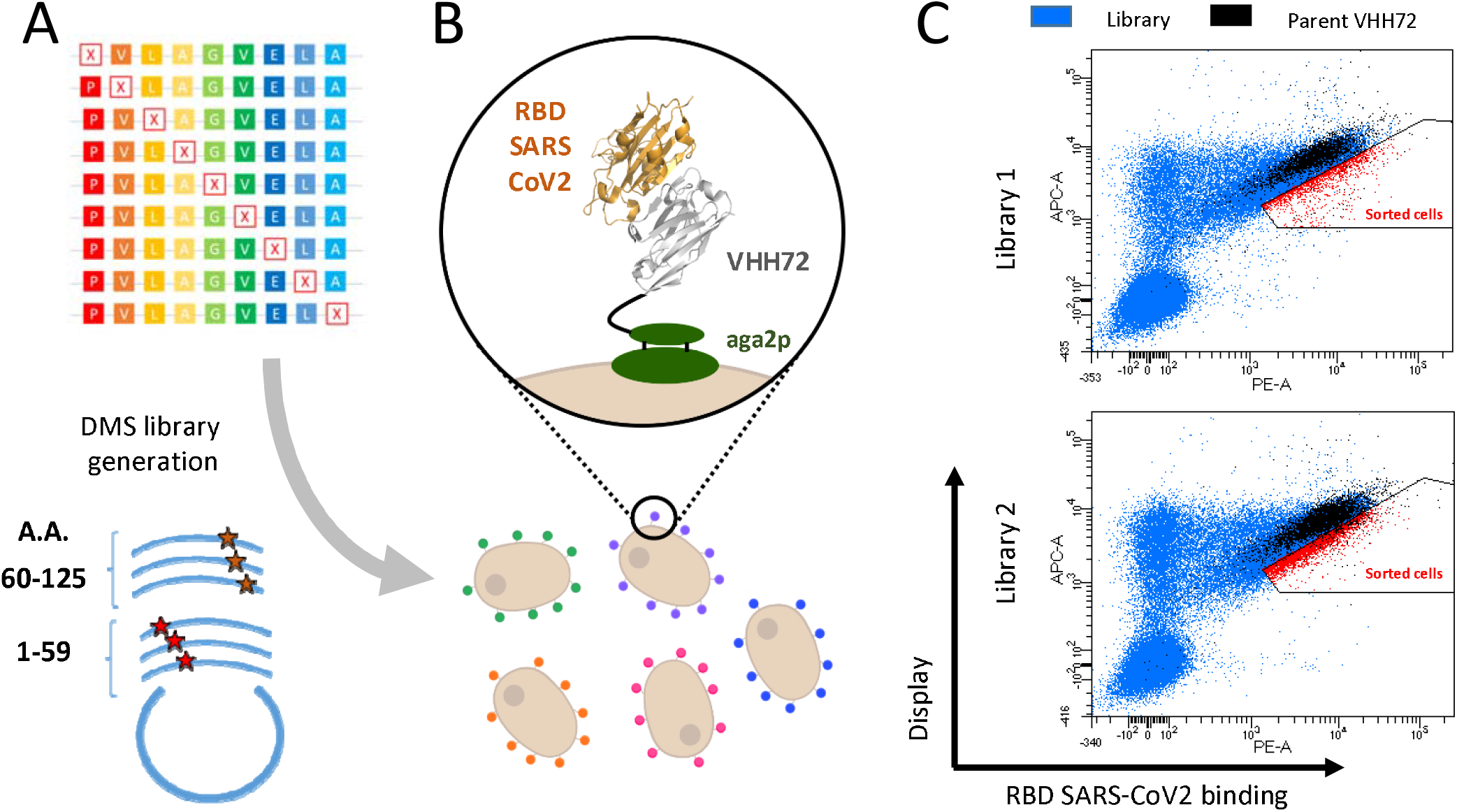
Deep Mutational Scanning probing VHH-72 binding to the RBD domain from SARS-CoV2 Spike protein. **(A)** Two DNA libraries of VHH72 harbouring a single mutation (each corresponding to regions encompassing amino acids 1-59 and 60-125 of the nanobody) were transformed into yeast using gap repair recombination. **(B)** General principle of functional screening by yeast surface display. Cells are incubated with biotinylated RBD antigen and labelled with secondary reporters before FACS analysis to determine VHH expression and antigen binding. **(C)** Bivariate flow cytometry analysis of libraries L1 and L2 of yeast cells expressing VHH72 variants on their surface. Cells were double-labelled with biotinylated antigen/Streptavidin–PE (RBD SARS-CoV2 binding) and anti-HA tag antibody coupled to APC (VHH expression). Cells corresponding to clones of the DMS libraries are represented in blue. Libraries were spiked with 10% of clonal cells expressing parental VHH72 along with egfp protein (represented in black) to discriminate cells with increased antigen binding levels. Selected cells (in red) were sorted and sequenced with Illumina Deep Sequencing.

For each library, plasmid DNA was sequenced for both sorted and unsorted cells to evaluate the respective frequency of clones in these populations. For each position and every substitution, enrichment ratios were calculated and represented as a functionality map (Fig 2 and Supp. Fig. 2).

**Fig. 2.**
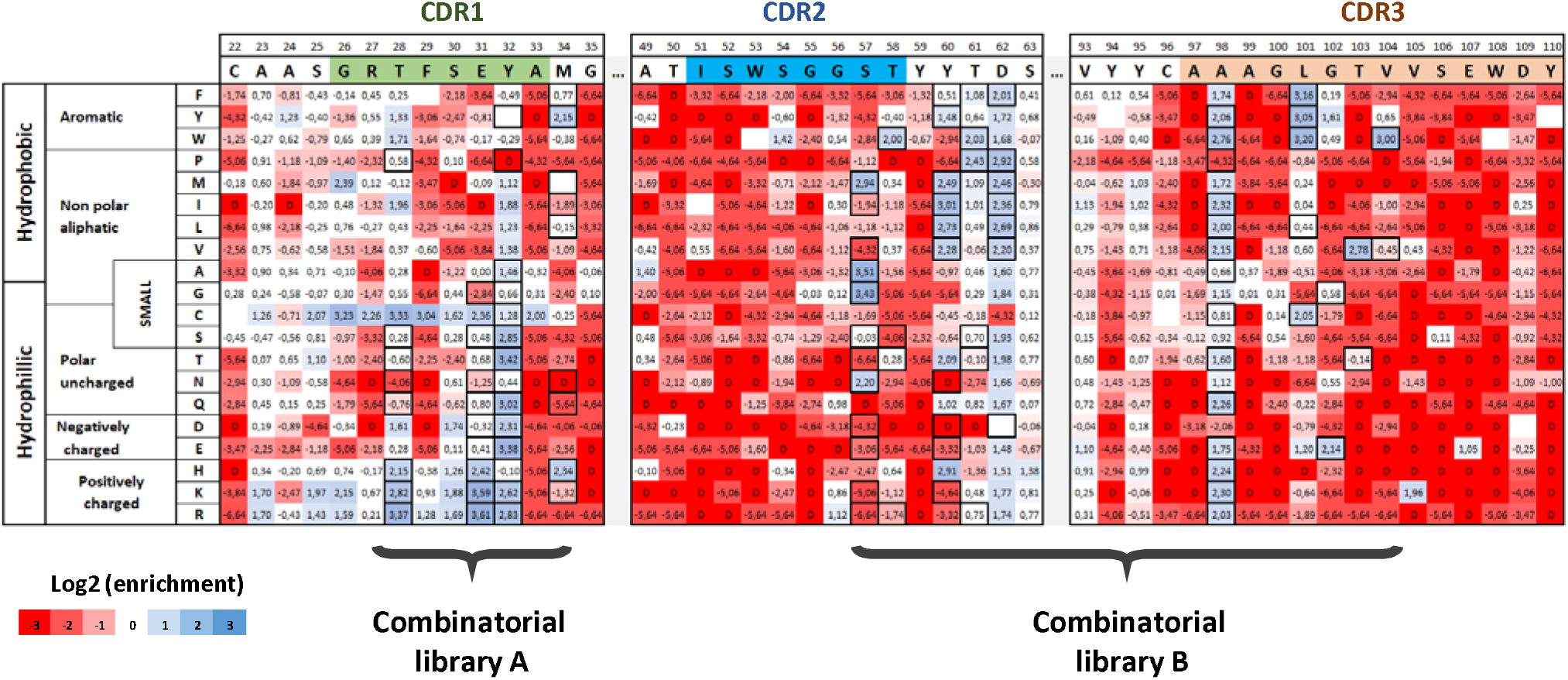
Paratope mapping of the VHH72 nanobody and design of combinatorial libraries for the selection of clones with improved affinity for SARS-COV-2 RBD antigen. NGS-based heatmap representing enrichment values of each VHH72 single mutant after functional sorting in FACS. Enrichment score is a base 2 log function of enrichment between sorted and unsorted VHH72 yeast populations for a given amino acid substitution. Corresponding table is coloured in blue for enriched mutations and in red for depleted mutations. Black squared substitutions were selected for the design of combinatorial libraries to identify VHH72 variants with improved binding properties. Due to the large generated diversity, two separate libraries were designed (with respective theoretical diversities of 4.14e4 and 2.63e7 clones).

Functionality map reveals uneven distributions of enriched clones (in blue) and depleted clones (in red). Most mutations in CDR2 and CDR3 are associated with a significant depletion in the sorted population, with the exception of few positions (e.g. A98 or L101). In contrast, several positions in CDR1 are relatively permissive (e.g. T28, E31 or Y32). More intriguingly, few positions demonstrate contrasted enrichment scores depending on the amino acid substitution involved, such as S57 for which four mutations are enriched (namely alanine, glycine, asparagine or methionine) while all charged and most hydrophobic amino acids have strong depletion scores. For positions T103 and V104, all substitutions have strong depletion scores except for respectively valine (T103) and tryptophan or tyrosine residues (V104) for which important enrichment is observed.

Based on these exhaustive mutational data, we generated optimized combinatorial libraries gathering all mutations with enrichment scores exceeding a threshold of 2 in the DMS experiment (i.e. a fourthfold enrichment). Custom degenerate primers were generated using the algorithm Swiftlib ^24^ and assembled in two libraries of approximately 4.10^4^ and 1.10^7^ clones, corresponding respectively to the CDR1 for library A and CDR2+CDR3 regions for library B. Both libraries were generated using splicing by overlap extension PCR (SOE-PCR). The parental amino acid was systematically included for all positions in the design, in contrast to cysteine substitution, which was excluded whenever possible. Because of the limited flexibility of codon degeneracy, a limited number of unwanted mutations were included in the libraries, without compromising their overall quality. The final design of the two libraries is indicated with black bolded squares (Fig 2).

After transformation in yeast cells, combinatorial libraries A and B were sorted to select variants with improved affinity for the SARS-CoV-2 antigen. Selection steps were initiated at 10 nM RBD SARS-CoV- 2, a concentration below the K_D_ of the parental antibody, to obtain a good dynamic range in antigen binding signal. Regardless of their overall good expression on the surface of yeast cells, best clones of library A demonstrated antigen binding levels similar to the parental clones, without visible affinity enhancement. In sharp contrast, library B comprises a large proportion of clones with improved RBD binding (approximately 30% of induced cells in the first selection round, Fig 3A). The top 2% clones (in the red gate) displayed a very significant increase in antigen binding. The corresponding cells were sorted and submitted to a second round of selection. In addition to the conventional selection of clones based on low antigen concentrations, we also introduced k_off_ based constraints to drive the selection of clones with long dissociation times. Thus, clones from the first round were incubated with 1 nM RBD biotinylated antigen for two hours, before washing and incubation with an excess amount of 100 nM non-biotinylated antigen to limit possible re-association of the antigen upon dissociation. After two more hours, clones with highest residual binding to biotinylated RBD were sorted (right panel, fig 3.A).

**Fig. 3.**
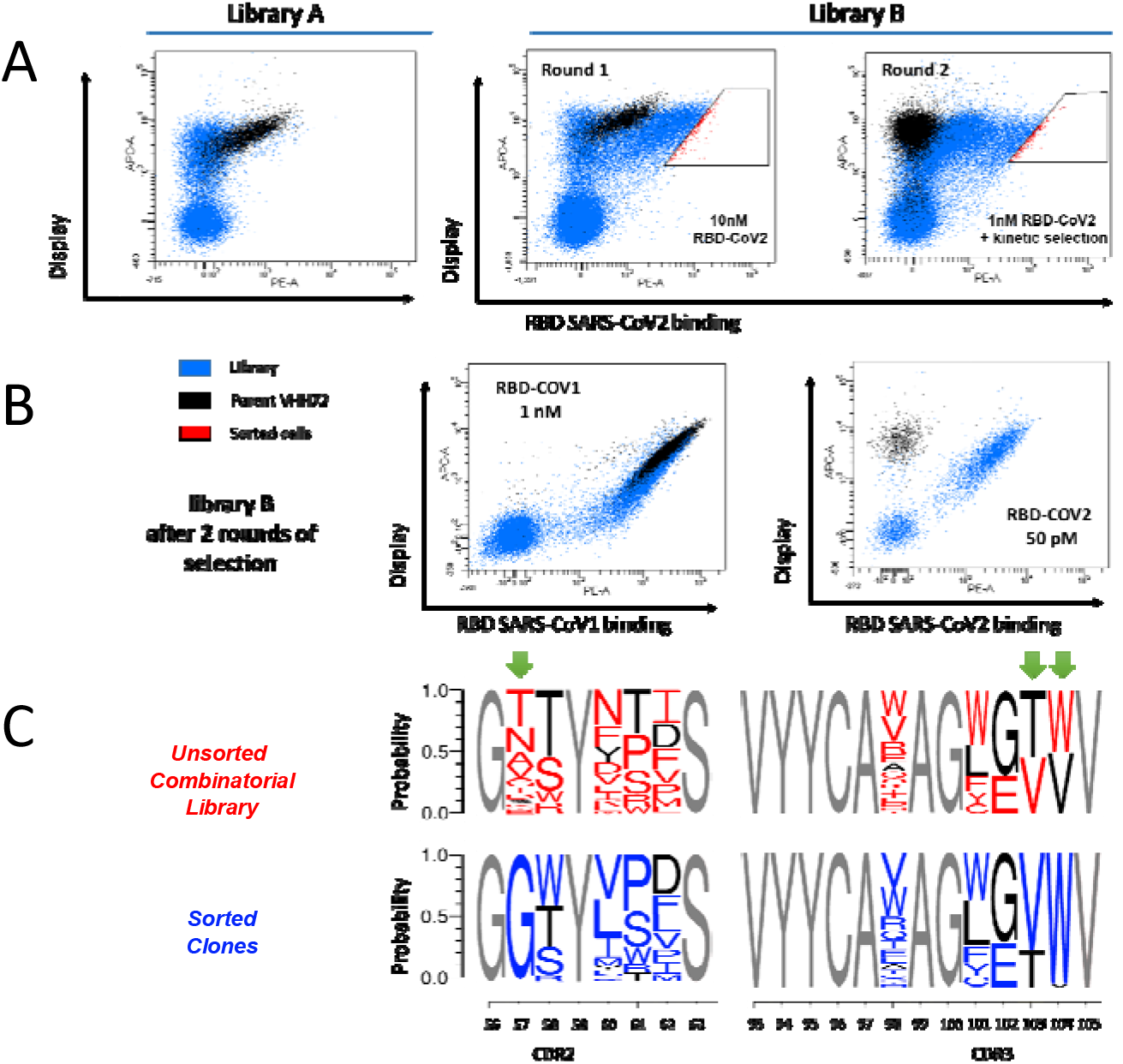
Yeast Surface Display based screening of libraries and sorting of VHH72 clones with improved binding for the RBD SARS-CoV-2 antigen. **(A)** Library A was incubated with 10nM biotinylated SARS-CoV-2 RBD and analysed in FACS. Few clones with improved antigen binding were detectable. Library B was sorted twice at respective concentrations of 10nM and 1nM biotinylated SARS-CoV-2 RBD. In the second round, a selection based on long dissociation times (slow k_off_) was performed. After equilibrium with 1nM biotinlylated antigen, an excess amount of 100 nM non-biotinylated antigen was introduced to drive dissociation and limit re-association. **(B)** Clones selected from library B after two rounds of FACS selection. All selected clones display strong binding to both SARS-CoV-1 and SARS-CoV-2 RBD antigens. **(C)** Sequence logos of clones contained in library B before sorting (in red) and after two rounds of FACS selection for improved affinity (in blue).

After these selection steps, enriched clones displayed very significant binding levels to SARS-CoV-2 RBD at low concentration of antigen (50 pM, fig. 3B), in contrast to the parental clone that exhibits negligible binding signal. Very interestingly, the vast majority of enriched clones still bind to SARS-CoV-1 RBD at 1nM (Fig. 3B). A second NGS campaign was undertaken to measure the evolution of amino acid frequencies in sorted and unsorted populations in the CDR2 and CDR3 regions. The frequency of mutants in the unsorted library is consistent with the theoretical library design (upper panel Fig. 3C). After two steps of selection, some mutations (in blue, Fig. 3C) are greatly enriched over the parental amino acid (in black), especially S57G, T103V and V104. Remarkably, S57G substitution is present in more than 97% of clones, replacing the eleven amino acids allowed in the library design. Similarly, tryptophan residue in position 104 (V104W) also has singular enrichment (96%) over valine, the parental amino acid. The amino acid distribution is altered for positions 58, 60, 61 and 62, with visible enrichment of hydrophobic residues over tyrosine at position 60 and threonine 61 being replaced principally by proline or serine residues.

To further investigate the gain of affinity conferred by these substitutions to these VHH molecules, we selected representative clones of highly enriched VHH based on the NGS data. We therefore chose five different VHH (variants VHH72.6, VHH72.65, VHH72.66, VHH72.71 and VHH72.76) each combining five to eight substitutions from the parental antibody (Sup. Data). Additionally, we examined the influence of the three principal substitutions, S57G, T103V and V104W, either separately or in combinations. Each mutant was expressed as a VHH-Fc construct. Upon transfection, VHH-Fc molecules were purified and their affinities for the different RBD antigens determined using BioLayer Interferometry (BLI). Each of the five engineered mutants exhibit strong sub-nanomolar affinities for RBD SARS-CoV-2 in a VHH-Fc format (Fig 4.A). The apparent antigen affinity for the Wuhan strain antigen is very high, with double-digit picomolar affinity constants for mutants VHH72.66 and VHH72.71. In addition, binding of these molecules to SARS-CoV-2 RBD is characterized by much longer dissociation rates compared to the parental molecule. For each VHH-Fc, the affinity is only marginally affected by mutations present within the different SARS-CoV-2 strains (Fig 4A). The five selected VHH-Fc molecules retain very good binding to the SARS-CoV-1 RBD domain. Their affinity constants are in the double-digit picomolar range, with a visible improvement over the parent molecule, whose affinity for the RBD domain of SARS-CoV-1 is in the nanomolar range.

**Fig 4.**
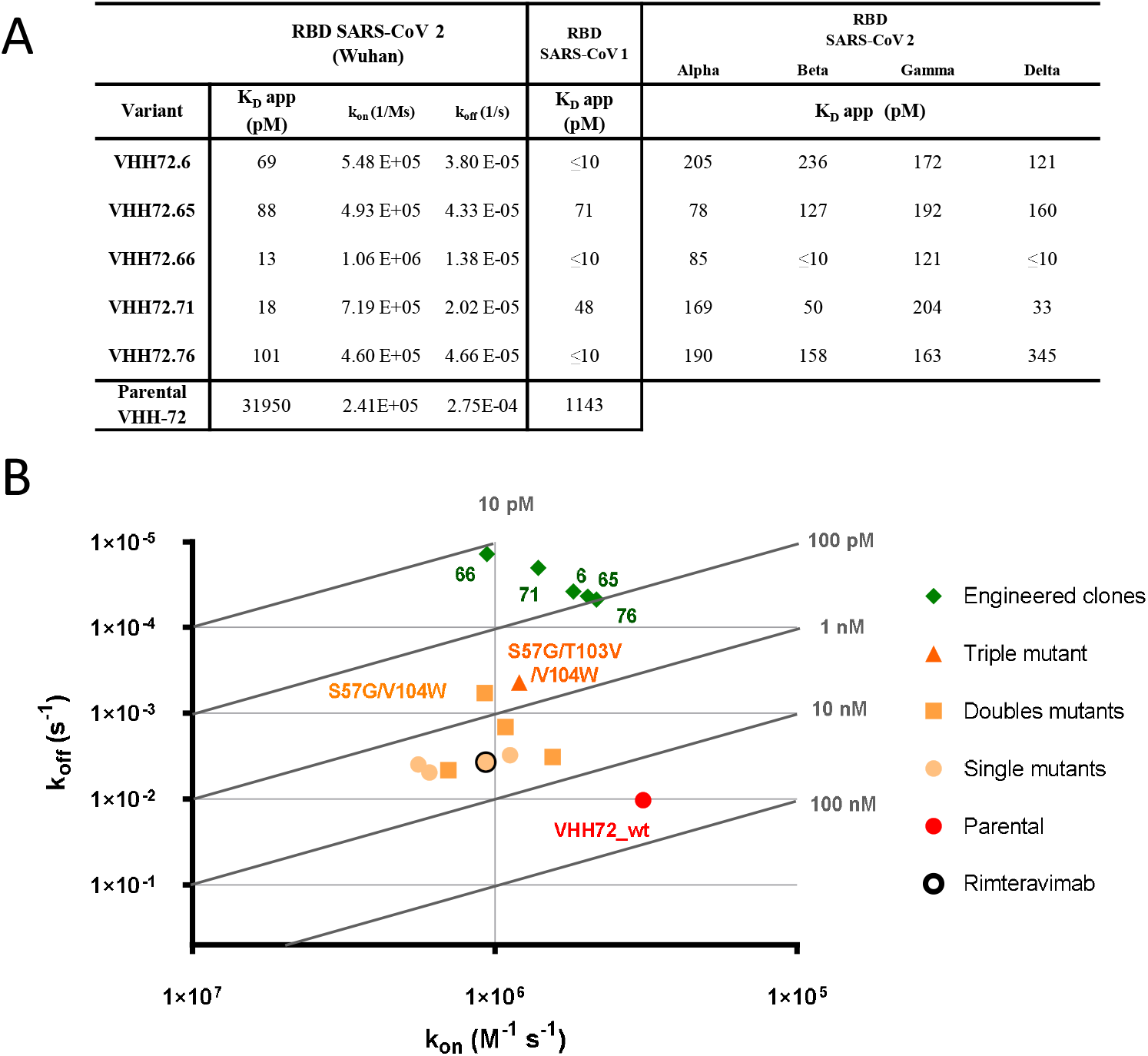
Affinity of VHH-Fc single-chain antibodies to RBD domains of SARS-CoV and SARS-CoV-2. **(A)** Bio-Layer Interferometry analysis of VHH-Fc immobilized proteins on anti-human Fc biosensors. Apparent binding kinetics of interaction between the VHH-Fc and the various RBD domains from SARS-CoV variants were evaluated in real time. Binding curves were fitted using a global 1:1 model. **(B)** Isoaffinity graph representation of k_on_ and k_off_ values for engineered clones and selected single, double and triple mutants, compared to the parental antibody and rimteravimab.

BLI experiments also showed that the S57G, T103V and V104W mutations individually and in combination increased the affinity for the SARS-COV-2 RBD antigen (in yellow, Fig 4B). Higher affinities were observed for the S57G/T103V/V104W triple mutant and for the S57G/V104W double mutation (in orange). These two combinations confer sub-nanomolar affinities to the VHH-Fc but are nonetheless significantly lower than those observed for the engineered mutants issued from the screening process (in green, Fig 4B). As expected given the constraints introduced during selection, the differences in apparent affinity constants observed are mainly explained by differences in dissociation constants. On the contrary, association constants were not significantly affected by the engineering steps. Overall, these data illustrate that these five engineered mutants not only show excellent affinities with long dissociation rates but also broad spectrum recognition of known SARS-CoV strains.

Next, we sought to verify that the engineered mutants were all able to antagonize the binding of RBD proteins to the human ACE2 receptor. To this end, we developed a competitive indirect ELISA test. Engineered VHH-Fc showed an increased ability to antagonize the binding of RBD to ACE2, with 50% inhibitory concentration (IC_50_) values lower than 0.1 μg/mL, up to 220-fold lower than those observed with the parent molecule VHH72-Fc (Fig. 5A and 5C). Our data show very similar IC_50_ values independently of the antigen tested (RBD of Wuhan, Beta or Delta strains) for each of the VHH-Fc molecules tested, confirming the large spectrum of recognition of those molecules.

**Fig 5.**
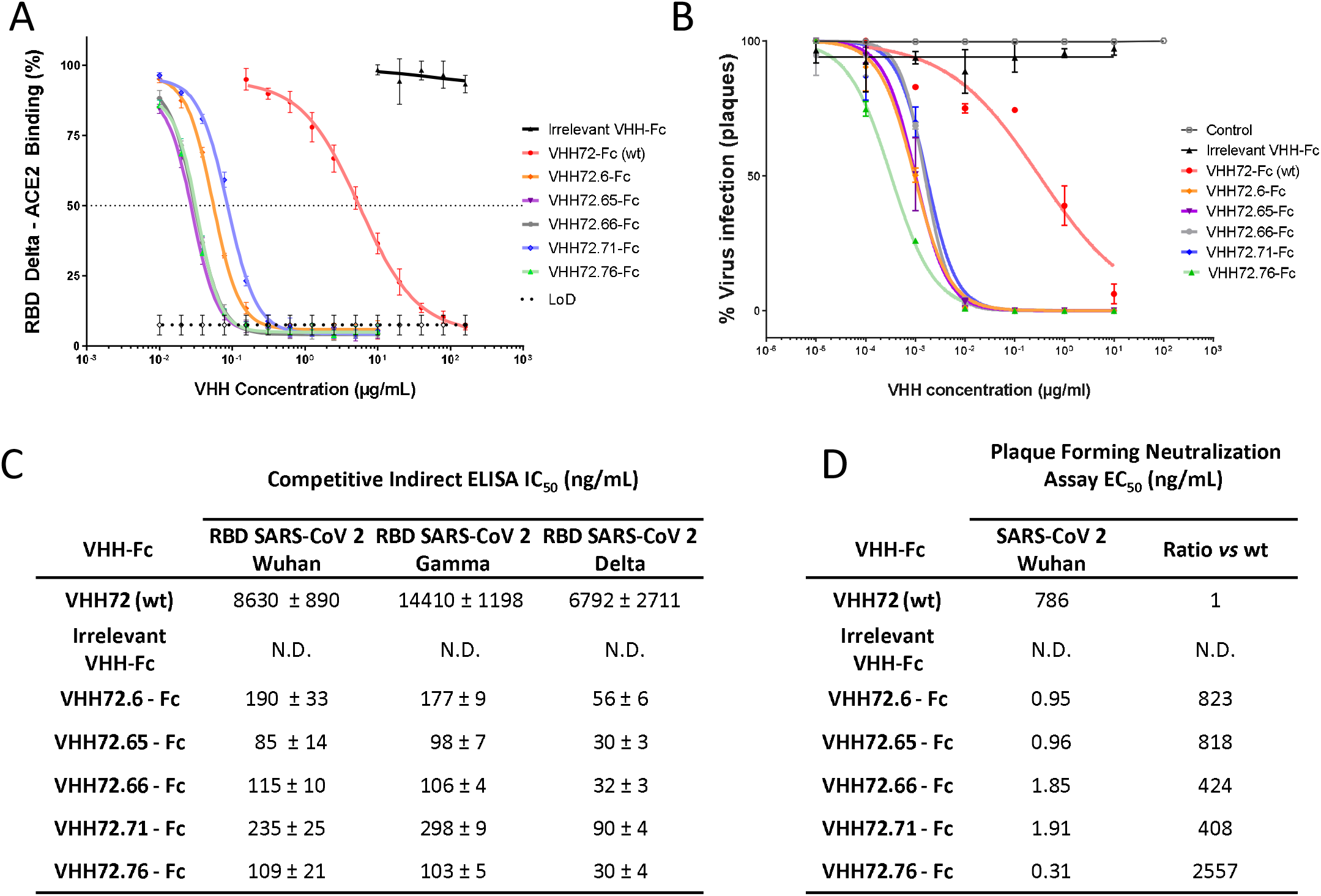
Protein-based competition ELISA and authentic virus cell-based neutralization assay. **(A)** Assessment of the ability of the selected VHH-Fc antibodies to block the interaction between ACE2 and the RBD domain of SARS-CoV-2 strain Delta in a competitive ELISA setup. Values represented correspond to three independent experiments. **(C)** Fifty percent inhibitory concentration (IC_50_) of the different VHH-Fc for the SARS-CoV2 variants Wuhan, Gamma and Delta by competitive ELISA was calculated. Data points represent mean values ± standard deviation of three independent experiments for each RBD from SARS-CoV2 variant. **(B) (D)** Neutralization of authentic SARS-CoV-2 virus (Wuhan strain) by the indicated VHH72-Fc constructs.

To confirm the ability of these single-chain antibodies to neutralize the SARS-CoV-2 virus and prevent its entry into human cells, we set up a cell-based virus neutralization assay. All VHH-Fc molecules tested demonstrate an excellent ability to neutralize cellular infection by SARS-CoV-2. The reference VHH (VHH72) has a neutralizing titer of 780 ng/mL (10 nM) and the optimized VHH-Fc all have neutralizing titers with improvements ranging from a factor of 408 for VHH72.71 to 2557 for VHH72.76. These results showed that engineered molecules exhibited very low neutralizing titers, down to 0.31 ng/mL (3.9 pM) for VHH72.76, objectively demonstrating the impact of sequence optimization.

We next tried to understand the mechanisms by which the introduced mutations in engineered VHH molecules increase the binding to the antigen. Based on the crystallographic structure of the VHH72/RBD SARS-CoV-1 complex, a model of the VHH72- SARS-CoV-2 RBD antigen was built and further refined by molecular dynamics simulation. Three independent 100 ns molecular dynamics trajectories were calculated. The initial model suggests that the VHH epitope (in cyan, Fig 6A) does not include any of the substitutions naturally present in the tested SARS-CoV-2 strains, which seems consistent with the observed cross-reactivity.

**Fig 6.**
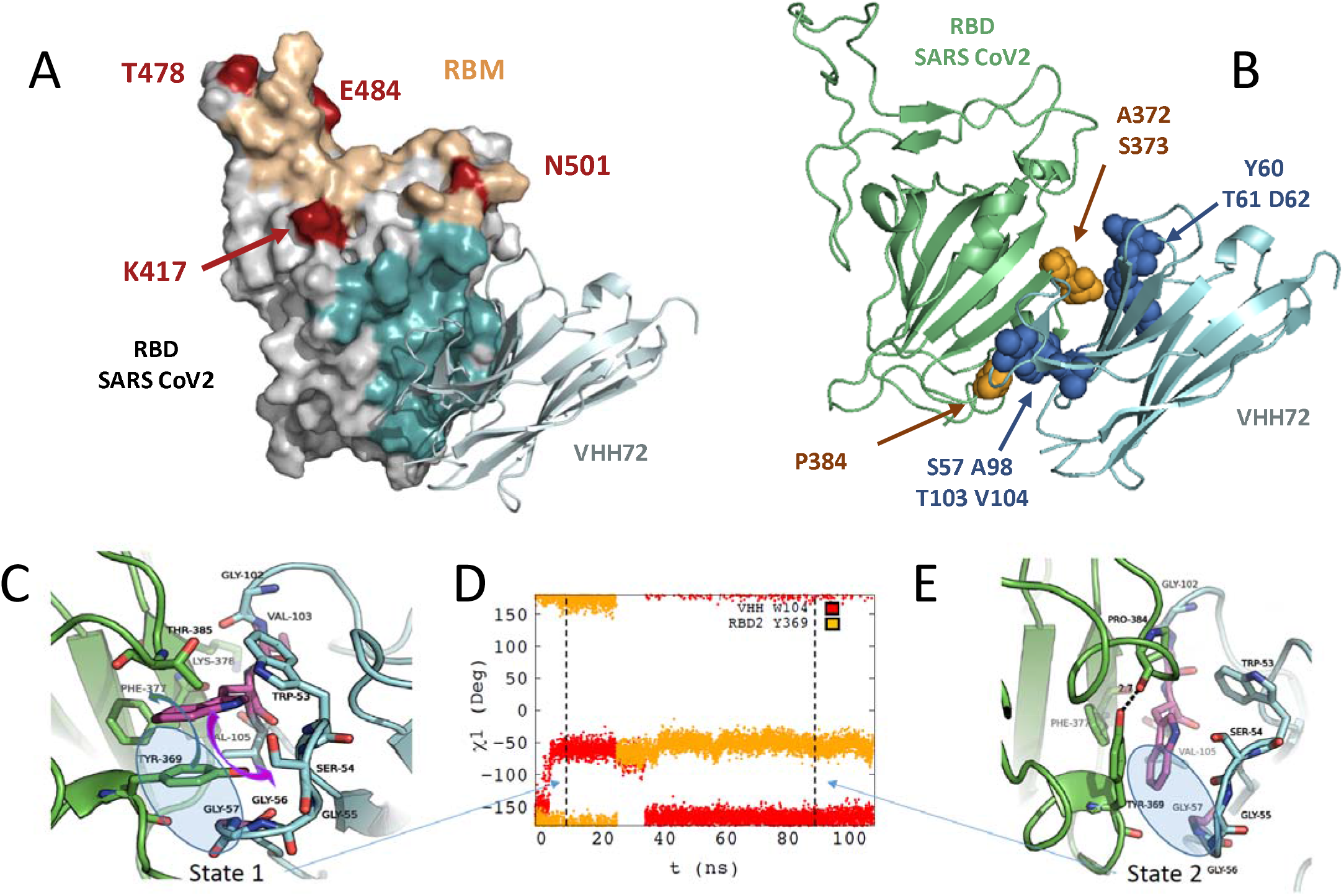
Molecular modelling of the interaction between VHH-72 harbouring substitutions S57G / T103V / V104W with SARS-CoV-2 RBD. **(A)** Homology model of interaction between VHH72 with substitutions S57G / T103V / V104W and SARS-CoV-2 RBD domain. Amino acid substitutions present in the different SARS-CoV-2 variants of concern (VOCs) are shown in red. RBD amino acids within 4.5Å of any VHH-72 atom were defined as the epitope and stained cyan. Residues included in the receptor binding motif (RBM) are shown in sand color. **(B)** Location of residues in the epitope of the VHH72 that diverge from SARS-CoV-2 and SARS-CoV-1 (in orange). Amino acid positions for which substitutions are present in our selected variants are shown in blue. (**C) (E)** Representative structures extracted from the MD simulation of the SARS-COV-2 RBD – VHH72 complex for state 1 **(C)** and state 2 **(E)**. Main chain of the proteins are shown in cartoon representation coloured in green for SARS CoV-2 RBD and in cyan for VHH72 with the exception of residues G57, V103 and W104 coloured in purple. Important residues for the interaction between the two proteins are shown in stick representation coloured by element. In Figure 6C, the movements of residues SARS-CoV-2 RBD Y369 and VHH72 W104 are shown using coloured arrows (green for SARS Cov2 RBD Y369 and purple for VHH72 W104). The χ1 monitored along the MD simulation are shown in **(D)** for residues W104 (in red) of VHH72 and Y369 (in orange).

The model shows an immediate proximity of the amino acid at position 57 to positions 103 and 104 of VHH72, which is consistent with the visible coevolution of these three residues in our data set. The model also suggests a direct interaction of the side chains of the amino acids at position 103 and 104 with the antigen, essentially by creating a network of van der Waals interactions with hydrophobic groups of SARS-CoV-2 RBD. Along the MD simulations, two states characterized by different χ1 values for residues W104 of VHH72 and Y369 of SARS-CoV-2 RBD are systematically found (Fig. 6E and Supp. Fig. 6A). In state 1, the side chain of VHH72 W104 is located in the center of a hydrophobic cavity formed by the side chains of F377 and Y369 of SARS-CoV-2 RBD and residue V105 and W53 of VHH72 (Figure 6C). The orientation of W104 is allowed because the side chain of SARS-CoV-2 RBD Y369 has moved from χ1 = -60° in SARS-CoV-1 RBD-VHH72 complex, structure 6WAQ) to χ1 = +180° in the VHH/SARS-CoV-2 model (State 2, Figure 6D and Supp. Fig 6A-C). It must be noted that the correlated changes of the χ1 dihedral angles of SARS-CoV-2 RBD Y369 and VHH72 W104 are accompanied by uncorrelated changes of their respective χ2 (Supp. Fig 6D-F). The S57G mutation is required to enable this χ1 rotation because the lack of a side chain in position 57 of VHH72 leaves space to accommodate the SARS-CoV-2 RBD Y369 side chain (Figure 6C, blue circle). In this regard, the distance between the CH2 atom of the side chain of VHH72 W104 and the Ca of VHH72 G57 clearly shows that a bulky side chain in position 57 could not accommodate a tryptophan residue in position 104 (Supp. Fig.7)). During MD trajectory 1, after about 25 ns, the side chains of SARS-CoV-2 RBD Y369 flip back to a position similar to the one observed in the SARS-CoV-1 RBD-VHH72 complex (χ1 = –60°, Figure 6E). This motion is accompanied by a rotation of the VHH72 W104 side chain, which also involves evolution of the χ2 of these two residues which reach an equilibrium state in the last part of the MD trajectories (Supp. Fig.6). Overall our modelling study explains the strong cooperativity between the VHH72 residues located at the interface with RBD antigens, in particular between W104 and G57 and to a lesser extent with V105 that contribute to form the hydrophobic cavity that accommodates the interface with SARS-CoV-2 RBD.

The mutations present in the engineered variants (variants VHH72.6, VHH72.65, VHH72.66 VHH72.71 and VHH72.76) are located in close proximity to the antigen (in blue, Fig 6B) and distributed in two distinct areas (S57, A98, T103, V104 and Y60, T61, D62 respectively). These mutations are present in the vicinity of three polymorphic amino acids that differ within the epitope of the RBD domains of SARS-CoV-1 and SARS-CoV-2 (in orange, Fig 6B). Positions S57, A98, T103, V104 are located in the environment of SARS-CoV-2 Proline 384 while positions Y60, T61 and D62 are in the proximity of SARS-CoV-2 A372 and S373. Engineered variants share common substitutions in the first region (specifically S57G, A98V, T103V and V104W) but incorporate different sets of substitutions in the second region. Hence, the slight differences in affinity and neutralization of the different variants can be attributed to substitutions at positions Y60, T61, and D62.

## Discussion

Monoclonal antibodies are becoming a major new class of drugs with very diverse fields of application, in oncology, inflammatory diseases but also for a number of severe infectious diseases such as infections by Respiratory Syncytial Virus ^25^, EBOLA ^26^ or SARS-CoV-2. Numerous methods are available to obtain new antibodies, whether by immunization of animals, screening of naive or synthetic libraries or by identification within the immune repertoires of convalescent patients for infectious diseases. In the context of the COVID19 pandemic, a wide range of approaches have been used to rapidly generate monoclonal antibodies against SARS-CoV-2 and to accelerate their clinical evaluation^27,28^. Over time, some of these antibodies have shown certain defects, in particular a decreased efficacy due to mutations present in emerging strains.^15,27^ These mutations alter not only their affinity, but also their ability to neutralize the virus and to provide an effective protection. As a result, a second generation of antibodies might be required to overcome these limitations and could be obtained by reshaping the initial sequences by conferring them the necessary properties, such as affinity and selectivity, to have optimal therapeutic efficacy for treatment in humans.

From this perspective, many teams have proposed a wide range of methods to generate candidates with the expected properties, mostly increased affinity.^29-32^ Affinity maturation aims at improving biological activity by adjusting the kinetic parameters of the binding to the target, which in turn may confer greater therapeutic efficacy.^25,33^ However, the magnitude of this effect depends largely on the epitope recognized by the antibody and the initial affinity along with the format of the antibody and its valence.^25^ In the context of the current COVID-19 pandemic, several studies have described affinity maturation of VHH or conventional antibodies to enhance their binding to SARS-CoV-2 antigens, by CDR-swapping approaches ^34^, saturation mutagenesis in CDRs ^35,36^ or light-chain shuffling^36^.

In recent years, Deep Mutational Scanning (DMS) approaches have emerged as a powerful tool for understanding protein/protein interactions. Deep Mutational Scanning (DMS) explores in a selected protein all possible unique substitutions, *i*.*e*. all unique mutations for each position. DMS defines mutational landscape of the protein and helps to understand the interaction modalities as recently shown for the RBD/ACE2 and RBD/antibodies interactions.^12,13,37,38^ In the present work, we used these systematic mutagenesis data to engineer new nanobodies with improved affinity and tailored selectivity. We named this process Deep Mutational Engineering (DME) and applied it to VHH72.

The therapeutic potential of VHH72 resides in its ability to neutralize SARS-CoV-2 in spite of its moderate affinity by recognizing an uncommon conserved class IV epitope^39^. The marked difference in affinity of the parent molecule for the antigens of the two viruses is probably explained by the presence of three substitutions found in the contact zones with VHH72 (residues A372, S373 and P384 in SARS-CoV-2 epitope are respectively threonine, phenylalanine and alanine residues in the SARS-CoV-1 epitope). On this basis, we first sought to improve the affinity of VHH72 by searching all individual favourable mutations conferring enhanced binding to SARS-CoV-2 RBD. After an exhaustive evaluation of the individual role of each amino acid and the influence of all possible substitutions, we demonstrated that it is possible to combine and cumulate individually favourable mutations in the VHH72 sequence. By generating combinatorial libraries and performing two consecutive selection steps, we were able to identify many mutants with very high affinity for the SARS-CoV-2 antigen using a Deep Mutational Engineering. After screening of variants for an increased binding to the SARS-CoV-2 antigen, we verified that the affinity for the initial antigen, the RBD domain of SARS-CoV- 1 was preserved. Interestingly, in the present case, selection of nanobodies for cross-reactivity was not required, as enriched clones had a preserved binding at low RBD SARS-CoV-1 concentrations. The best clones characterized even showed a significantly improved affinity for the SARS-CoV-1 antigen, below 100 pM. Otherwise, it would have been perfectly conceivable to introduce an additional selection step to select clones based on their SARS-CoV-1 RBD binding in order to retain only clones with the desired cross-reactivity.

The vast majority of our selected clones incorporate a subset of mutations consisting of S57G, T103V and V104W substitutions. This combination alone confers a significant gain in affinity for the SARS-CoV-2 antigen, by approximately 2 logs, resulting in a sub-nanomolar molecule. The DMS data for these three key positions show very low permissiveness, with very few favourable substitutions. This contrasting pattern seems ultimately more promising in terms of affinity maturation than the positions with many enriched substitutions (e.g. D62 or A98) which could indicate a certain permissiveness or an influence on the expression levels. Interestingly, all the molecules emerging from the screening incorporate other substitutions (between two to five additional changes) that further increase their affinity for the antigen, up to two-digit picomolar affinities. This suggests that beyond this subset of core mutations, additional substitutions contribute to further increase the affinity for the antigen. The accumulation of several favourable and potentially additive changes could be one of the important advantages of this DMS-based affinity maturation method. This could potentially allow for greater affinity gains especially in comparison with random mutagenesis approaches such as error-prone PCR methods ^29^. Indeed, our method generates libraries with a large proportion of active clones whereas accumulation of several random mutations per clone would most probably result in mostly inactive molecules.

After engineering, our selected VHH-Fc molecules are not only capable of recognising both SARS-CoV-1 and SARS-CoV-2 antigens, but also emerging SARS-CoV-2 variants with very high affinity and balanced affinity profiles. It seems possible that the selected constructs will be less sensitive to future mutations than many other antibodies, for two reasons. First, the epitope has a high degree of conservation between SARS-CoV-1 and SARS-Cov-2 viruses, which differ by three positions in this epitope, only. This high degree of conservation within sarbecoviruses could be explained by the apparent role of this region in folding and conformational changes of the spike protein^18^. Moreover, antibodies produced in the sera of patients infected by SARS-CoV-1 or SARS-CoV-2 induce a weak level of cross-reactivity suggesting that the two viruses only share few epitopes.^19,21^ The VHH72 epitope is not among the dominant epitopes in the antibody repertoire of infected or vaccinated individuals and could even be qualified as subdominant. The humoral response would be expected to exert limited selective pressure on this region and therefore mutations in the VHH72 epitope would not provide any evolutionary advantage for the virus to escape an immune response.

Thus, our VHH-Fc seems to be one of the few described high affinity and broad-spectrum molecules, such as sotrovimab^17^, ADG2^22^ or S2H97^5^ antibodies. Importantly, the gains in affinity observed for the engineered molecules translate into an increase in their potential to not only neutralize the interaction between RBD and the human ACE2, but also the entry of the SARS-CoV-2 virus into human cells. Unlike many class IV antibodies which often are weakly neutralizing,^40^ our molecules demonstrate a strong capacity to neutralize the SARS-CoV-2 virus, with low EC50 in the ng/mL and picomolar range. Recently, rimteravimab, which is derived from VHH72 and incorporates a single substitution corresponding to S56A, has shown protection in mice and Syrian hamster animal models.^23^ This molecule is currently in human clinical trials but has an affinity in the nanomolar range, evaluated at 6 nM,^23^ that is about 100 to 200 times lower than our best candidates and a significantly lower neutralization potential. Thus, it would be interesting to verify that the gain of affinity acquired by molecular engineering translates into a better protection in animals either for therapeutic applications as a curative treatment.

Mutations conferring an increase of the affinity for the SARS-CoV-2 antigen appear to be in close proximity to the epitope. In particular, T103V and V104W are located in the direct vicinity of the polymorphic amino acids, including position P384, thus contributing to a better accommodation of the paratope to the SARS-CoV-2 antigen. Altogether, our results suggest that it is possible to tailor antibodies with known specificity to other related antigens, provided that the original antibody or nanobody has somehow a detectable initial binding to the antigen. In the case of an emergence of new strains of existing pathogens, some of the available antibodies could thus serve as templates for generating new effective antibodies of high affinity and custom specificity.

Overall, thanks to their broad recognition spectrum, their high affinity and their ability to neutralize the SARS-CoV-2 virus, these broadly neutralizing antibodies (bnAbs) could be molecules of interest for clinical use in the fight against the current SARS-CoV-2 pandemic.

## Material and Methods

### DMS Library design and generation of combinatorial libraries for affinity maturation

Two DMS libraries of VHH72 variants with single amino acid mutations were generated by splicing by overlap extension PCR (SOE-PCR) using degenerate NNK primers. A library was generated for the two targeted regions (positions 1 to 59 and 61 to 125) corresponding respectively to a diversity of 1888 and 2112 DNA variants.

Design of mutagenic primers containing degenerate codon was performed using SwiftLib (http://rosettadesign.med.unc.edu/SwiftLib/). Two libraries were designed corresponding respectively to the CDRH1 and CDRH2+CDRH3 regions.

### Yeast surface display of VHH

Preparation of competent yeast cells EBY100 (ATCC® MYA-4941) and library transformation were performed according to Benatuil *et al* ^41^. Gap repair transformations were made in plasmid pNT VHH72 between restriction sites NheI and NotI with 1 μg of digested vector and a molar ratio of 12:1 (library/digested vector). After transformation, cell were cultivated in 250 mL of SD-CAA medium ^42^. After a passage to an OD_600_ of 0.25, cells were grown at 30°C until OD_600_ 0.5-1.0 and re-suspended in 50 mL of SG-CAA for induction^42^.

### Flow cytometry

For library sorting, 10^7^ to 2.10^8^ induced cells were washed PBSF (PBS, BSA 0.1%) and resuspended in PBSF containing the appropriate antigen concentration using an appropriate volume to avoid ligand depletion as performed in Hunter *et al*. ^43^. After 3 hours incubation at 20°C with agitation, cells were washed with ice-cold PBSF to avoid dissociation. Cells were incubated on ice for 15 minutes with anti- HA antibody (Invitrogen HA Tag Mouse anti-Tag, DyLight® 650 conjugate, Clone: 2-2.2.14; 1:100 dilution) and Streptavidin-PE (Thermo Fisher scientific; catalog number S866; 1:100 dilution). Cells were subsequently washed with ice-cold PBSF and sorted with a BD FACS Aria™ III cytometer using BD FACSdiva™ software.

### Deep sequencing and analysis of NGS data

Plasmid DNA of each yeast population was extracted using Zymoprep Yeast Plasmid Miniprep II and prepared for sequencing as described in Medina-Cucurella and Whitehead ^44^. Two-step PCR was performed to amplify the region of interest and add Illumina adapters and barcodes for multiplexing. Deep sequencing was performed with an Illumina ISeq 100 device (2×150 bp, 300 cycles) with at least 300,000 reads per population. Reads were demultiplexed and each sample was processed separately using the Galaxy platform (https://usegalaxy.org/) using the functions described in Blankenberg *et al* ^45^. First, paired reads were joined (Fastq Joiner). A trim was then performed (Fastq Trimmer) on reads to keep just the region of interest in the correct frame. A quality filter (Filter FASTQ) was applied to eliminate reads with a minimum quality score under 30. Next, DNA sequences were translated in protein sequences and identical sequences were grouped. Sequences not repeated at least two times were filtered out. Using the software RStudio, single-mutants were selected to allow calculation of enrichment ratios for each single mutation.

### Production and purification of the VHH72-Fc ligands

VHH72-Fc and RBD SARS-CoV-2 constructs were obtained by transient transfection of HEK293 Freestyle™ (Thermo). Synthetic genes corresponding to selected VHH72 mutants were ordered to IDT as gblocks and cloned into the mammalian expression vector pcDNA 3.4 VHH72-Fc. Genes coding for the various RBD domains were cloned in the pCAGGS RBD-SARS-CoV-2 plasmid, which was a kind gift from Florian Krammer lab. HEK293 Freestyle™ were transiently transfected at a density of 2.5 10^6^ cells/mL in 100mL Freestyle medium (Thermo-Fisher) by addition of 150 μg plasmid and 1.8 mL of linear polyethylenimine (PEI, 0.5 mg/ml) (Polysciences). After 24h, 100 mL of fresh medium were added. After seven days at 37°C, 120 rpm, 8% CO2, supernatant was purified using HiTrap Protein A for VHH-Fc constructs or HisTrap Excel for RBD constructs, following the manufacturer’s instructions (GE Healthcare). Size-exclusion chromatography was performed (HiPrep Sephacryl-S-200HR or S100HR) with PBS.

### Affinity measurement by BioLayer Interferometry

Binding kinetics were determined using an Octet RED96 instrument (ForteBio). Anti-hIgG Fc Capture (AHC) Biosensors (Fortebio) were loaded with VHH-Fc molecules (50 nM) for 60 seconds. After baseline using kinetic buffer (PBS, BSA 0.5% (w/v) and Tween 20 0.05% (v/v)), association of RBD SARS-CoV-2 or RBD SARS-CoV-1 (Sinobiological) was measured at different concentrations (50 nM to 0.78 nM) for 300 seconds before dissociation in kinetic buffer. Data of the control without antigen were subtracted from all binding curves and binding kinetics were fitted using a global 1:1 Langmuir-binding model.

### Virus neutralization Assay

VERO E6 (ATCC CRL-1586) cells are seeded in DMEM medium (Gibco) + 5% fetal calf serum (FCS) (Hyclone) in sterile 12-well plates (Falcon) at 2.5 10^5^ cells per well and incubated for 24 hours at 37°C - 9% CO_2_. Purified SARS-CoV-2 virus (strain 2019-nCov/Italy-INMI1, provided by the European Virus Archive goes Global, EVAg) is diluted in DMEM + 2.5 % FCS to 400 PFU/mL and mixed 1:1 with each VHH dilution, ranging from 200 μg/mL to 0.2 ng/mL. They are incubated under agitation for one hour at 37°C - 9 % CO_2_. Then, the culture medium of each plate containing the VERO E6 cells is removed and 500 μL of each VHH/virus mixture is added to each well in duplicate. The virus suspension is incubated for 45 minutes at 37°C - 9 % CO_2_ and then removed. Two milliliters of carboxymethyl cellulose (Merck) 2 % in DMEM + 10 % FCS are then added. After 72 hours of incubation at 37°C 9% CO_2_, the medium is removed and the cells are stained with crystal violet for 20 minutes at room temperature. After removal of the crystal violet and washing in phosphate-buffered saline, the lysis plaques in each well are counted.

### Molecular modelling of the VHH72-SARS-CoV-2 RBD

We used the structure of SARS-CoV-2 RBD-VHHE as template to build a model of the SARS-CoV-2 RBD-VHH72 complex. The initial structure of VHH72 was built by homology modelling using the MODELLER software ^46^ and the coordinates of VHH72 in PDB structure 6WAQ as template ^18^. The coordinates of the SARS-CoV-2 RBD were taken from the structure 7KN5 ^47^ and the model of VHH72 was positioned by superimposing the model of VHH72 on the VHHU ones. This initial model was refined by a protocol of energy minimisation and molecular dynamics simulation under positional restraints. During this first step, the solvent was treated implicitly. Then, the resulting structure of the SARS-CoV-2 RBD-VHH72 was studied by molecular dynamics in explicit solvent. All calculations were carried out with NAMD ^48^ with the CHARMM36 force field ^49^. The initial structure was immersed in a TIP3P water box using the solvate tool of VMD ^50^. The limit of the box was set such that any solute atom was located at a distance at least equal to 12 Å of the limit in each direction. Then, the system was neutralized using the autoionize plugin of VMD with Na^+^ ions. The two steps protocol MD simulations (equilibration and production) was achieved in the NPT ensemble using Langevin dynamics. A time step of 2fs was used in the equilibrium and the production runs with the SHAKE method to constrain bond vibration involving hydrogens^51^. A cutoff of 12 Å was used for non-bonded interactions and a dielectric constant of 1.0 for electrostatic interactions. Multiple timestepping was used with the rRESPA method to calculate long range interaction forces every 4 fs. We used periodic boundary conditions and Particle Mesh Ewald (PME) method ^52^ to treat the long-range electrostatics interactions with a real space grid of 1Å. The temperature was set to 310K and the pressure at 1.0 atm. The first step started with several 3000 steps cycles of energy minimization under positional restraints. From one cycle to the next one, the force constant applied for the positional restraints were gradually decreased to reach 0 kcal. mol^-1^ A^-2^ at the end of the minimization step. Then, a 1ns equilibration step of MD simulation was run after random initialisation of the velocities. This step was followed by a 110 ns MD production step. The random initialisation of the velocities allow to run independent trajectories. To ensure reproducibility of the results, three 100 ns MD trajectories of the SARS-CoV-2 RBD-VHHE were calculated starting with different initial velocities. Analysis of the trajectories was achieved using in-house scripts written in the macrolanguage of CHARMM v42b1^53^. Figures were produced with PyMol [PyMOL Molecular Graphics System, Version 2.0 Schrödinger, LLC.] and Gnuplot 5.1.

### Competitive indirect enzyme linked immunoassay (competitive ELISA)

Nunc™ MaxiSorp 96-well Immuno-Plates (Thermo Fisher Scientific, Illkirch, France) were coated with 200 μL/well of AffinityPure goat anti-mouse IgG+IgM (H+L) antibody (Jackson Immuno Research Laboratories Inc., Pennsylvania, USA) at 10 μg/mL in 50 mM potassium phosphate buffer, and incubated overnight (16h) at 22 ± 2 °C. Plates were then saturated by addition of 300 μL per well of enzyme immunoassay (EIA) buffer (0.1M phosphate buffer, pH 7.4, 0.15 M NaCl, 0.1 % bovine serum albumin, and 1 % sodium azide) and stored at 4 °C until use (at least for 24 h). Two-fold serial dilutions of the different VHH-Fc were prepared in EIA buffer ranging from 0.029 μg/mL to 30 μg/mL (for VHH72.6-Fc, VHH72.65-Fc, VHH72.66-Fc, VHH72.71-Fc, VHH72.76-Fc) or from 0.470 to 480 μg/mL (for VHH72-Fc and irrelevant VHH-Fc). Simultaneously, a solution of Human ACE2-mouse Fc tag (Sino Biological Inc., Eschborn, Germany) at 900 ng/mL in EIA buffer and a solution of biotinylated RBD (recombinant RBD from the SARS-CoV-2 variants Wuhan, Gamma and Delta) at 300 ng/mL in EIA buffer were prepared. Solutions were distributed in the coated plates as follows: 50 μL/well of ACE- 2-mouse Fc tagged, 50 μL/well of the different concentrations of each VHH-Fc (or EIA buffer in the control wells to determine the B0 (100%) binding signal), and 50 μL/well of biotinylated RBD. Plates were incubated at 4 °C for 18h, and, after three washes, 150 μL of Pierce™ Streptavidin-PolyHRP (ThermoFisher Scientific, Illkirch, France) diluted 25000 fold in EIA buffer without sodium azide, were distributed in each well and incubated for 30 minutes at 22 ± 2 °C. Finally, plates were extensively washed, 150 μL of TMB (1-Strep Ultra ™B-ELISA, ThermoFisher Scientific, Illkirch, France) distributed in each well, and after 30 minutes incubation at 22 ± 2 °C, 150 μL of a 2N H_2_SO_4_ solution were added to stop the enzymatic reaction. Absorbances at 450 and 620 nm were determined. Results were expressed as percentage of B0 (the signal obtained in the absence of VHH-Fc (control wells)).

## Acknowledgements

This research was funded by the French joint ministerial program of R&D against CBRNE threats. This publication was supported by the European Virus Archive goes Global (EVAg) project that has received funding from the European Union’ s Horizon 2020 research and innovation programme under grant agreement No 653316.

## References

1 Weinreich, D. M. et al. REGN-COV2, a Neutralizing Antibody Cocktail, in Outpatients with Covid-19. N. Engl. J. Med. 384, 238–251, doi:10.1056/NEJMoa2035002 (2021).

2 Libster, R. et al. Early High-Titer Plasma Therapy to Prevent Severe Covid-19 in Older Adults. N. Engl. J. Med. 384, 610–618, doi:10.1056/NEJMoa2033700 (2021).

3 Chen, P. et al. SARS-CoV-2 Neutralizing Antibody LY-CoV555 in Outpatients with Covid-19. N. Engl. J. Med. 384, 229–237, doi:10.1056/NEJMoa2029849 (2021).

4 Raybould, M. I. J., Kovaltsuk, A., Marks, C. & Deane, C. M. CoV-AbDab: the coronavirus antibody database. Bioinformatics 37, 734–735, doi:10.1093/bioinformatics/btaa739 %J Bioinformatics (2020).

5 Starr, T. N. et al. SARS-CoV-2 RBD antibodies that maximize breadth and resistance to escape. Nature 597, 97–102, doi:10.1038/s41586-021-03807-6 (2021).

6 Weisblum, Y. et al. Escape from neutralizing antibodies by SARS-CoV-2 spike protein variants. Elife 9, doi:10.7554/eLife.61312 (2020).

7 Chen, R. E. et al. Resistance of SARS-CoV-2 variants to neutralization by monoclonal and serum-derived polyclonal antibodies. Nat. Med. 27, 717–726, doi:10.1038/s41591-021-01294-w (2021).

8 Wang, P. et al. Antibody resistance of SARS-CoV-2 variants B.1.351 and B.1.1.7. Nature 593, 130–135, doi:10.1038/s41586-021-03398-2 (2021).

9 Liu, Z. et al. Identification of SARS-CoV-2 spike mutations that attenuate monoclonal and serum antibody neutralization. Cell Host Microbe 29, 477–488.e474, doi:10.1016/j.chom.2021.01.014 (2021).

10 Kistler, K. E. & Bedford, T. Evidence for adaptive evolution in the receptor-binding domain of seasonal coronaviruses OC43 and 229e. eLife 10, e64509, doi:10.7554/eLife.64509 (2021).

11 Eguia, R. T. et al. A human coronavirus evolves antigenically to escape antibody immunity. PLoS pathogens 17, e1009453, doi:10.1371/journal.ppat.1009453 (2021).

12 Starr, T. N., Greaney, A. J., Dingens, A. S. & Bloom, J. D. Complete map of SARS-CoV-2 RBD mutations that escape the monoclonal antibody LY-CoV555 and its cocktail with LY-CoV016. Cell Rep Med 2, 100255, doi:10.1016/j.xcrm.2021.100255 (2021).

13 Greaney, A. J. et al. Comprehensive mapping of mutations in the SARS-CoV-2 receptor-binding domain that affect recognition by polyclonal human plasma antibodies. Cell Host & Microbe 29, 463–476.e466, doi:https://doi.org/10.1016/j.chom.2021.02.003 (2021).

14 Widera, M. et al. Bamlanivimab does not neutralize two SARS-CoV-2 variants carrying E484K in vitro. medRxiv, 2021.2002.2024.21252372, doi:10.1101/2021.02.24.21252372 (2021).

15 Planas, D. et al. Reduced sensitivity of SARS-CoV-2 variant Delta to antibody neutralization. Nature 596, 276–280, doi:10.1038/s41586-021-03777-9 (2021).

16 Lv, H. et al. Cross-reactive Antibody Response between SARS-CoV-2 and SARS-CoV Infections. Cell Rep 31, 107725, doi:10.1016/j.celrep.2020.107725 (2020).

17 Pinto, D. et al. Cross-neutralization of SARS-CoV-2 by a human monoclonal SARS-CoV antibody. Nature 583, 290–295, doi:10.1038/s41586-020-2349-y (2020).

18 Wrapp, D. et al. Structural Basis for Potent Neutralization of Betacoronaviruses by Single-Domain Camelid Antibodies. Cell 181, 1004–1015 e1015, doi:10.1016/j.cell.2020.04.031 (2020).

19 Ju, B. et al. Human neutralizing antibodies elicited by SARS-CoV-2 infection. Nature 584, 115–119, doi:10.1038/s41586-020-2380-z (2020).

20 Bates, T. A. et al. Cross-reactivity of SARS-CoV structural protein antibodies against SARS-CoV-2. Cell Rep 34, 108737, doi:https://doi.org/10.1016/j.celrep.2021.108737 (2021).

21 Wang, C. et al. A human monoclonal antibody blocking SARS-CoV-2 infection. Nature communications 11, 2251, doi:10.1038/s41467-020-16256-y (2020).

22 Rappazzo, C. G. et al. Broad and potent activity against SARS-like viruses by an engineered human monoclonal antibody. Science 371, 823, doi:10.1126/science.abf4830 (2021).

23 Schepens, B. et al. An affinity-enhanced, broadly neutralizing heavy chain-only antibody protects against SARS-CoV-2 infection in animal models. Sci Transl Med, eabi7826, doi:10.1126/scitranslmed.abi7826 (2021).

24 Jacobs, T. M., Yumerefendi, H., Kuhlman, B. & Leaver-Fay, A. SwiftLib: rapid degenerate-codon-library optimization through dynamic programming. Nucleic Acids Res. 43, e34, doi:10.1093/nar/gku1323 (2015).

25 Wu, H. et al. Ultra-potent Antibodies Against Respiratory Syncytial Virus: Effects of Binding Kinetics and Binding Valence on Viral Neutralization. J. Mol. Biol. 350, 126–144, doi:https://doi.org/10.1016/j.jmb.2005.04.049 (2005).

26 Corti, D. et al. Protective monotherapy against lethal Ebola virus infection by a potently neutralizing antibody. 351, 1339–1342, doi:doi:10.1126/science.aad5224 (2016).

27 Nathan, R. et al. A Narrative Review of the Clinical Practicalities of Bamlanivimab and Etesevimab Antibody Therapies for SARS-CoV-2. Infectious Diseases and Therapy 10, 1933–1947, doi:10.1007/s40121-021-00515-6 (2021).

28 Davide, C., Lisa, A. P., Gyorgy, S. & David, V. Tackling COVID-19 with neutralizing monoclonal antibodies. Cell, doi:10.1016/j.cell.2021.05.005 (2021).

29 Simons, J. F. et al. Affinity maturation of antibodies by combinatorial codon mutagenesis versus error-prone PCR. mAbs 12, 1803646, doi:10.1080/19420862.2020.1803646 (2020).

30 Packer, M. S. & Liu, D. R. Methods for the directed evolution of proteins. Nat. Rev. Genet. 16, 379–394, doi:10.1038/nrg3927 (2015).

31 Rajpal, A. et al. A general method for greatly improving the affinity of antibodies by using combinatorial libraries. Proc. Natl. Acad. Sci. U. S. A. 102, 8466–8471, doi:10.1073/pnas.0503543102 (2005).

32 Skamaki, K. et al. In vitro evolution of antibody affinity via insertional scanning mutagenesis of an entire antibody variable region. 117, 27307–27318, doi:10.1073/pnas.2002954117 %J Proceedings of the National Academy of Sciences (2020).

33 Maynard, J. A. et al. Protection against anthrax toxin by recombinant antibody fragments correlates with antigen affinity. Nat. Biotechnol. 20, 597–601, doi:10.1038/nbt0602-597 (2002).

34 Zupancic, J. M. et al. Directed evolution of potent neutralizing nanobodies against SARS-CoV-2 using CDR-swapping mutagenesis. Cell Chem Biol, doi:10.1016/j.chembiol.2021.05.019 (2021).

35 Schoof, M. et al. An ultrapotent synthetic nanobody neutralizes SARS-CoV-2 by stabilizing inactive Spike. Science 370, 1473–1479, doi:10.1126/science.abe3255 (2020).

36 Rouet, R. et al. Potent SARS-CoV-2 binding and neutralization through maturation of iconic SARS-CoV-1 antibodies. mAbs 13, 1922134, doi:10.1080/19420862.2021.1922134 (2021).

37 Starr, T. N. et al. Deep Mutational Scanning of SARS-CoV-2 Receptor Binding Domain Reveals Constraints on Folding and ACE2 Binding. Cell 182, 1295–1310 e1220, doi:10.1016/j.cell.2020.08.012 (2020).

38 Greaney, A. J. et al. Mapping mutations to the SARS-CoV-2 RBD that escape binding by different classes of antibodies. Nature communications 12, 4196–4196, doi:10.1038/s41467-021-24435-8 (2021).

39 Zupancic, J. M. et al. Engineered Multivalent Nanobodies Potently and Broadly Neutralize SARS-CoV-2 Variants. Advanced Therapeutics 4, 2100099, doi:https://doi.org/10.1002/adtp.202100099 (2021).

40 Shrestha, L. B., Tedla, N. & Bull, R. A. Broadly-Neutralizing Antibodies Against Emerging SARS-CoV-2 Variants. 12, doi:10.3389/fimmu.2021.752003 (2021).

41 Benatuil, L., Perez, J. M., Belk, J. & Hsieh, C. M. An improved yeast transformation method for the generation of very large human antibody libraries. Protein Eng. Des. Sel. 23, 155–159, doi:10.1093/protein/gzq002 (2010).

42 Sivelle, C. et al. Fab is the most efficient format to express functional antibodies by yeast surface display. mAbs, 1–10, doi:10.1080/19420862.2018.1468952 (2018).

43 Hunter, S. A. & Cochran, J. R. Cell-Binding Assays for Determining the Affinity of Protein-Protein Interactions: Technologies and Considerations. Methods Enzymol. 580, 21–44, doi:10.1016/bs.mie.2016.05.002 (2016).

44 Medina-Cucurella, A. V. & Whitehead, T. A. Characterizing Protein-Protein Interactions Using Deep Sequencing Coupled to Yeast Surface Display. Methods Mol. Biol. 1764, 101–121, doi:10.1007/978-1-4939-7759-8_7 (2018).

45 Blankenberg, D. et al. Manipulation of FASTQ data with Galaxy. Bioinformatics 26, 1783–1785, doi:10.1093/bioinformatics/btq281 (2010).

46 Šali, A. & Blundell, T. L. Comparative Protein Modelling by Satisfaction of Spatial Restraints. J. Mol. Biol. 234, 779–815, doi:https://doi.org/10.1006/jmbi.1993.1626 (1993).

47 Koenig, P. A. et al. Structure-guided multivalent nanobodies block SARS-CoV-2 infection and suppress mutational escape. Science 371, doi:10.1126/science.abe6230 (2021).

48 Phillips, J. C. et al. Scalable molecular dynamics on CPU and GPU architectures with NAMD. J Chem Phys 153, 044130, doi:10.1063/5.0014475 (2020).

49 Huang, J. & MacKerell, A. D., Jr. CHARMM36 all-atom additive protein force field: validation based on comparison to NMR data. J Comput Chem 34, 2135–2145, doi:10.1002/jcc.23354 (2013).

50 Humphrey, W., Dalke, A. & Schulten, K. VMD: visual molecular dynamics. J Mol Graph 14, 33–38, 27-38, doi:10.1016/0263-7855(96)00018-5 (1996).

51 Ryckaert, J. P., Ciccotti, G. & Berendsen, H. J. C. Numerical-Integration of Cartesian Equations of Motion of a System with Constraints - Molecular-Dynamics of N-Alkanes. J Comput Phys 23, 327–341 (1977).

52 Darden, T., York, D. & Pedersen, L. Particle Mesh Ewald - an N.Log(N) Method for Ewald Sums in Large Systems. J Chem Phys 98, 10089–10092 (1993).

53 Brooks, B. R. et al. CHARMM: the biomolecular simulation program. J Comput Chem 30, 1545–1614, doi:10.1002/jcc.21287 (2009).

