## Supplemental Data for "Deep Mutational Engineering of broadly-neutralizing and picomolar affinity nanobodies to accommodate SARS-CoV-1 & 2 antigenic polymorphism"

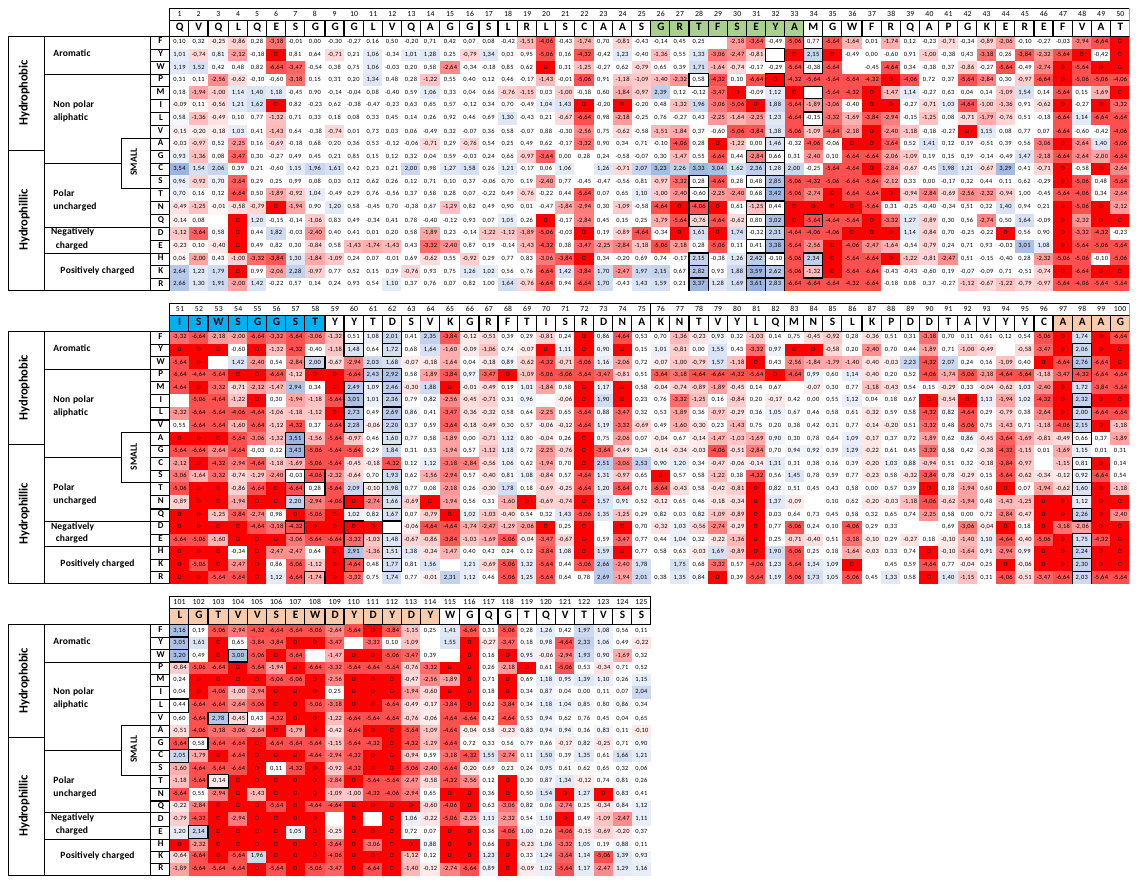

**CDR1**

**CDR2**

**CDR3**

AA selected for combinatorial libraries

**Supplementary Figure 1:** NGS-based heatmap representing enrichment values of each VHH72 single mutant after functional sorting in FACS. Enrichment score is a base 2 log function of enrichment between sorted and unsorted VHH72 yeast populations for a given amino acid substitution. Blue is for enriched mutations and red is for depleted mutations. Black bolded squares represent the final design of the two libraries.

**Supplementary Figure 1**

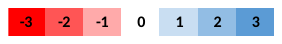

**Log2 (enrichment)**

**Supplementary Figure 2**

**Library 1 (CDR1) Diversity (DNA) 4,147e+4 clones**

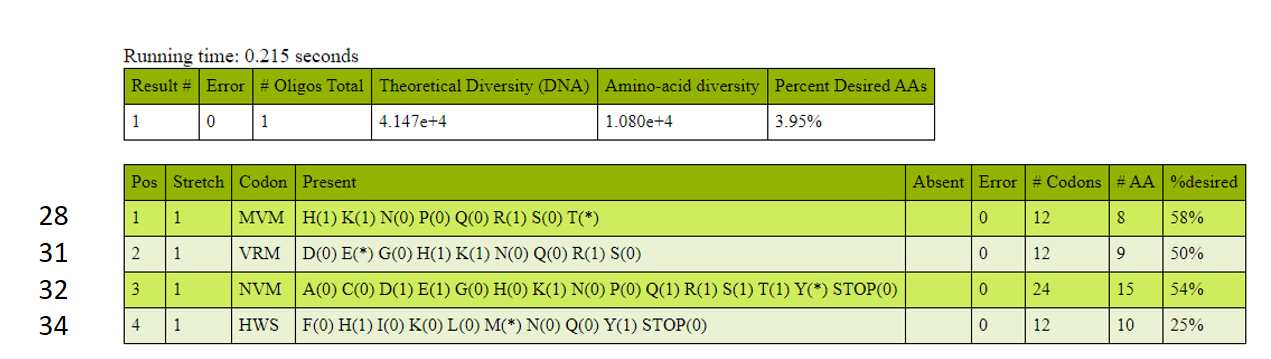

AA Position

**Library 2 (CDR2 + CDR3) Diversity (DNA) 2.63e+7 clones**

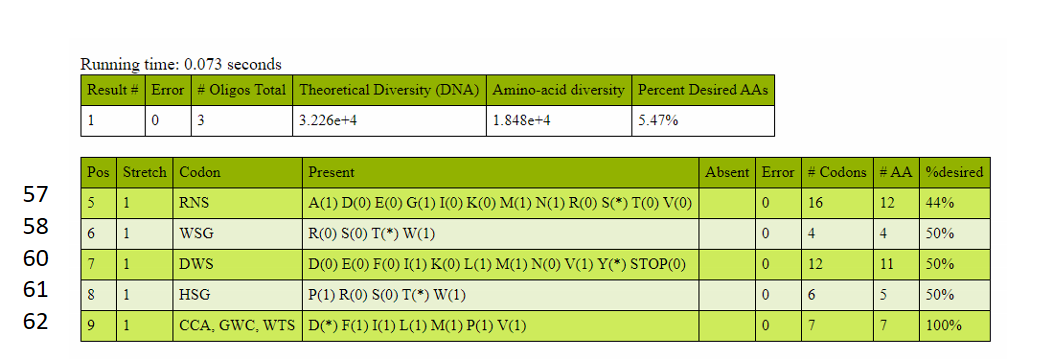

AA Position

AA Position

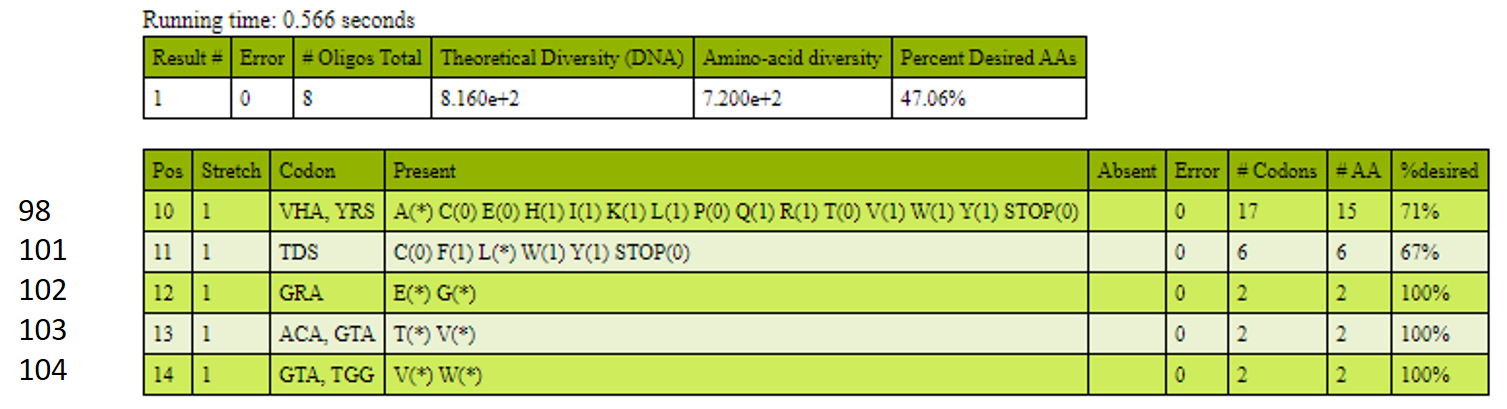

**Supplementary Figure 2: Optimized combinatorial libraries generation :** Based on DMS data, custom degenerate codon primers were generated using the algorithm Swiftlib and assembled in two libraries of approximately 4x104 and 2x107 clones, corresponding respectively to the CDR1 position 28-31-32-34 for library A and CDR2+CDR3 position 57-58-60-61-62; 98-101-102-103-104 for library B.

**Supplementary Figure 3**

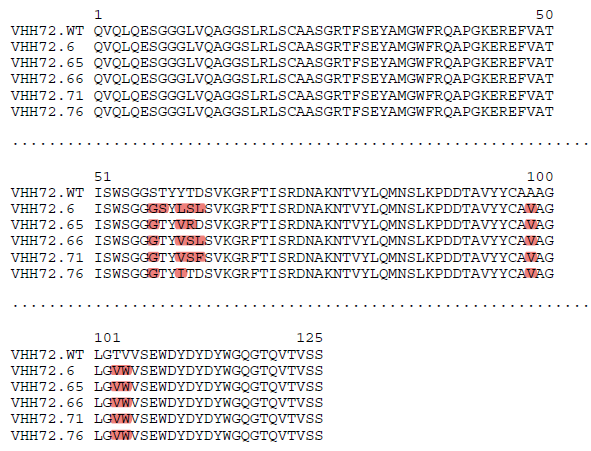

Mismatch

**Supplementary Figure 3: Multiple protein alignment:** Multiple protein alignment of VHH72.WT and 5 highly enriched VHH based on the NGS data. Red colour represents mismatch between VHH.

**Supplementary Figure 4A**

| **Mutants** |  |  | **RBD SARS-CoV 2 Variant of Interest** | | | |
| --- | --- | --- | --- | --- | --- | --- |
|  | **RBD SARS-CoV 1** | **RBD SARS-CoV 2** | **Alpha** | **Beta** | **Gamma** | **Delta** |
| **6** | 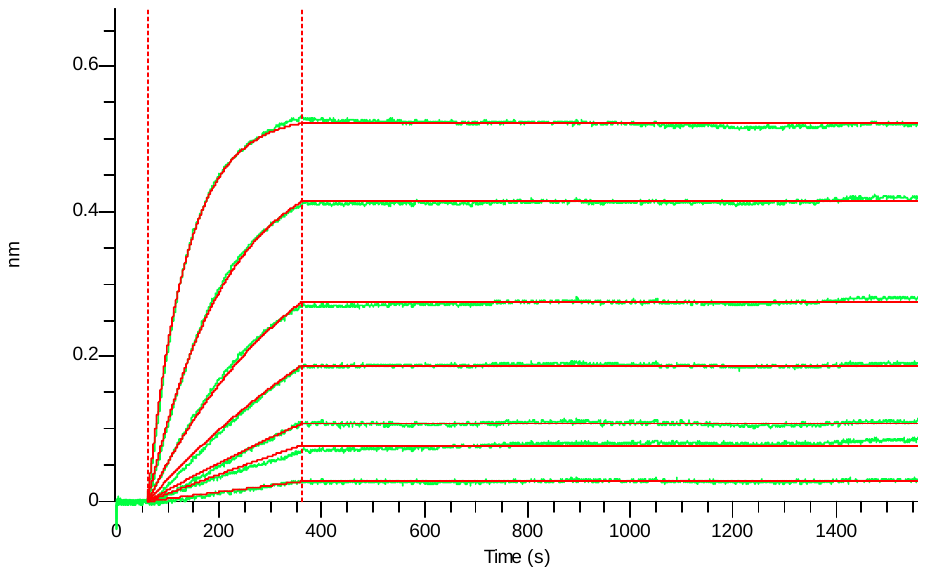 | 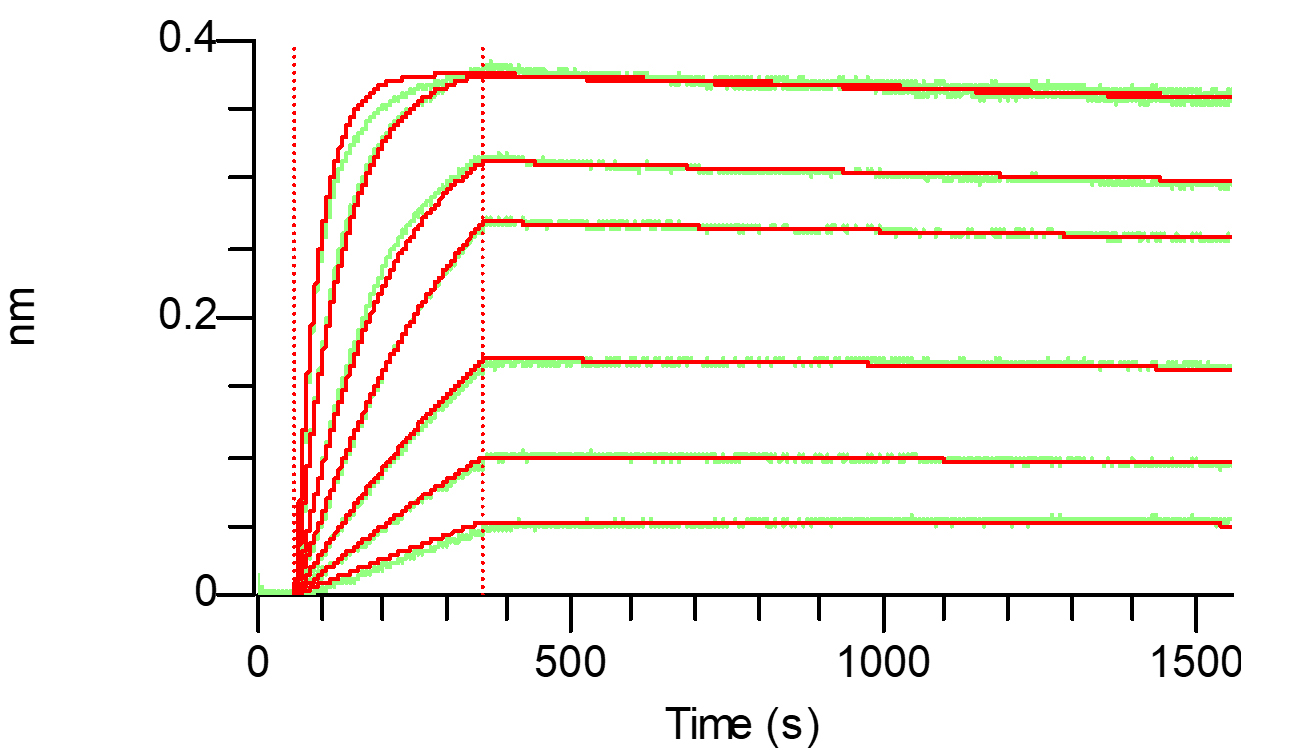 | 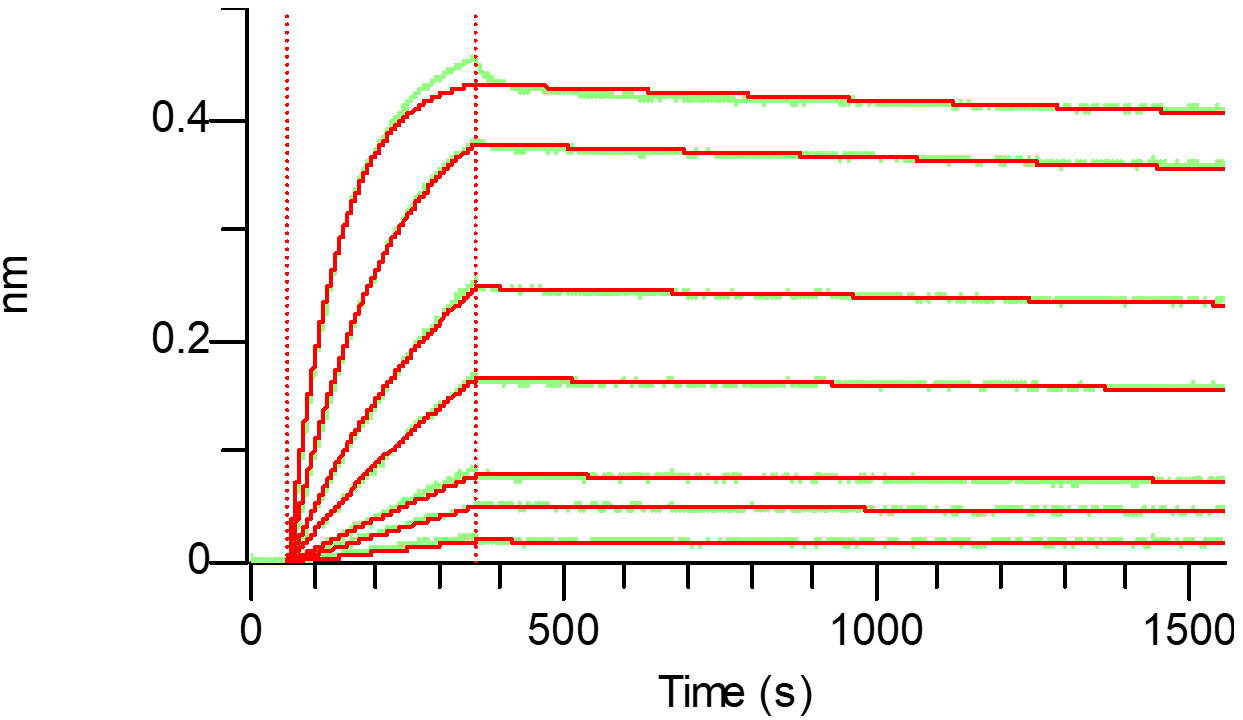 | 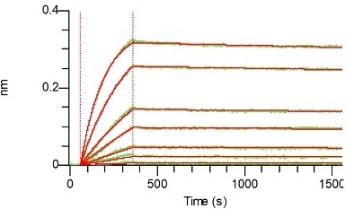 | 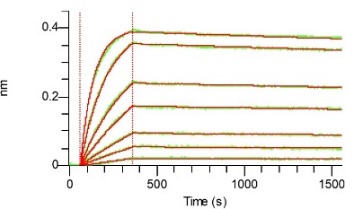 | 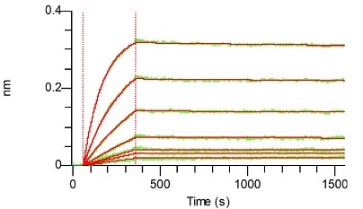 |
| **65** | 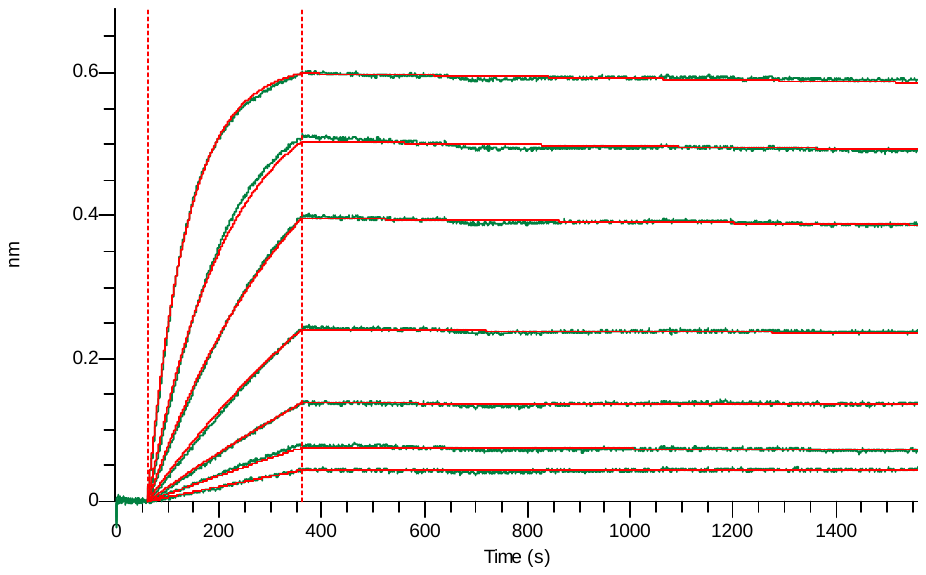 | 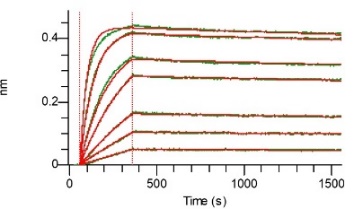 | 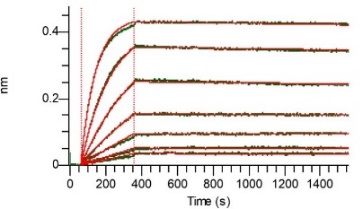 | 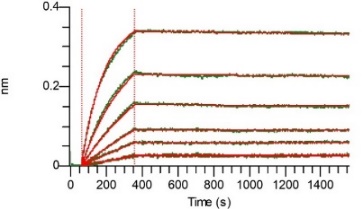 | 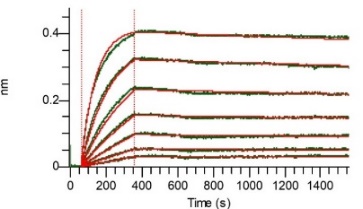 | 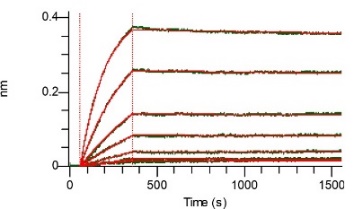 |
| **66** | 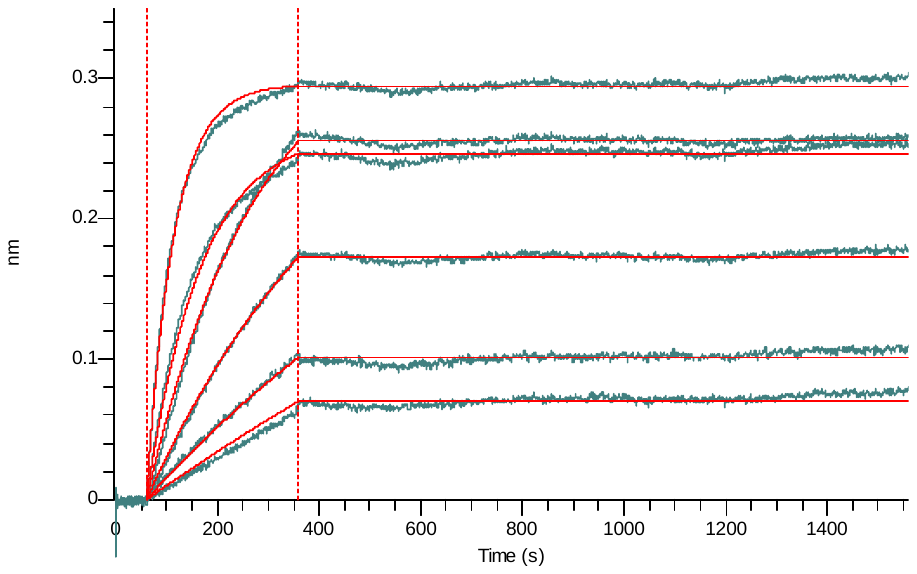 | 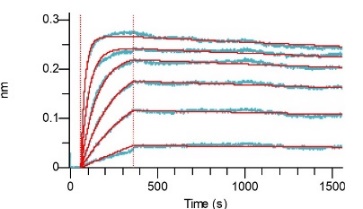 | 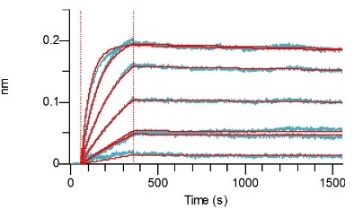 | 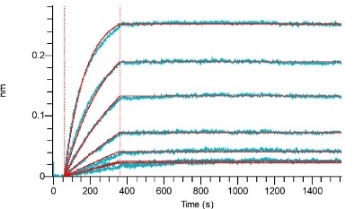 | 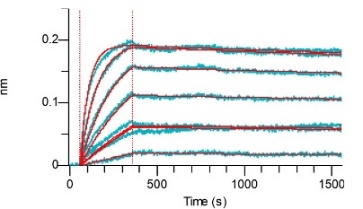 | 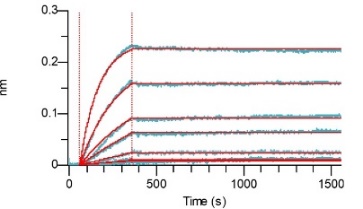 |
| **71** | 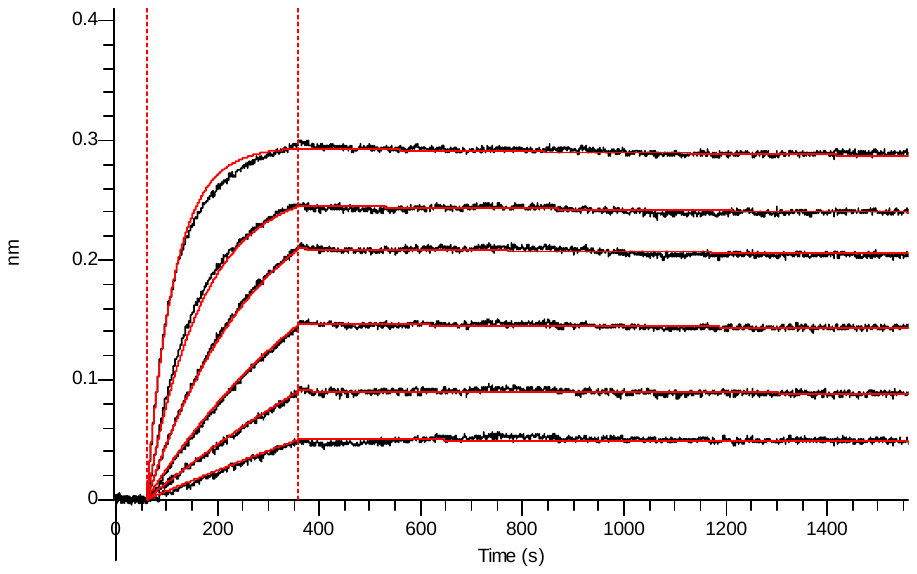 | 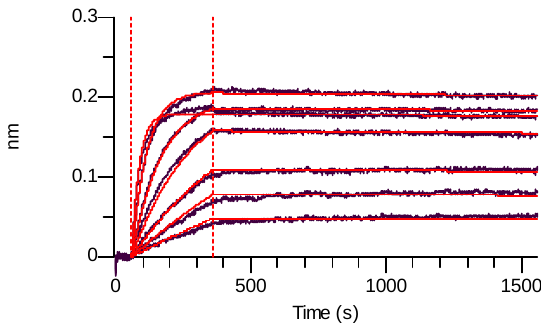 | 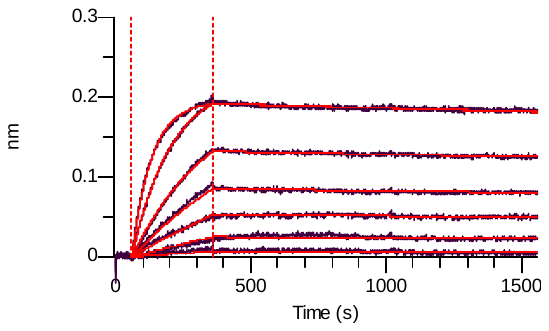 | 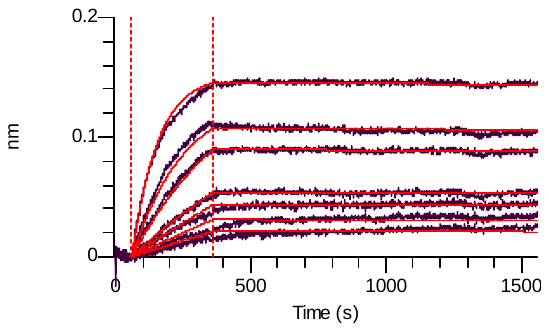 | 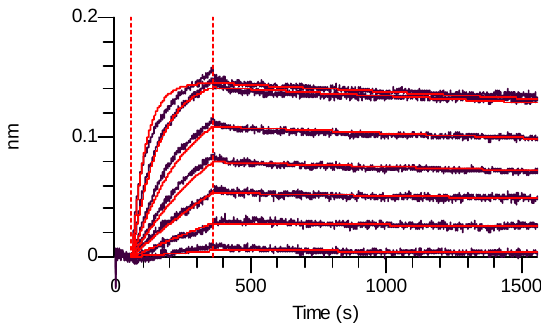 | 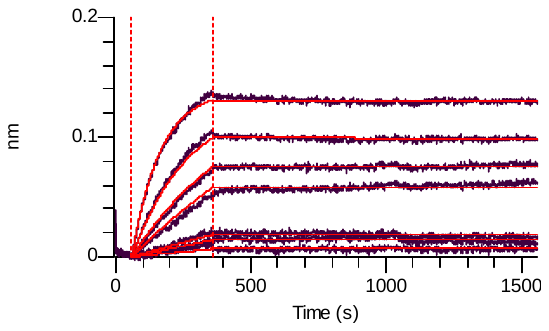 |
| **76** | 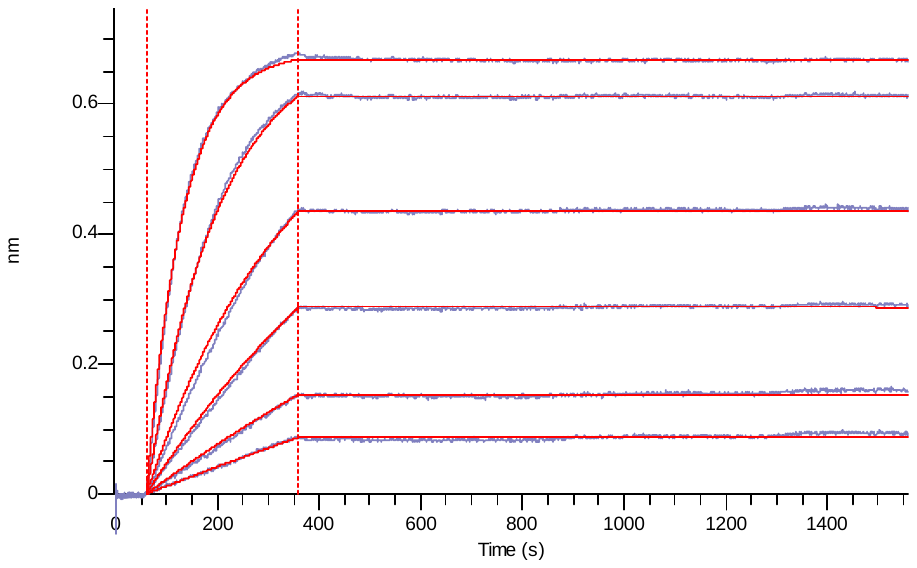 | 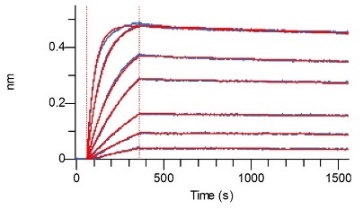 | 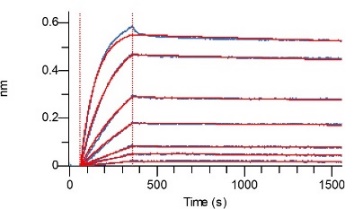 | 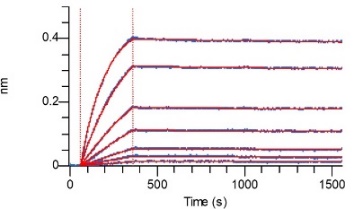 | 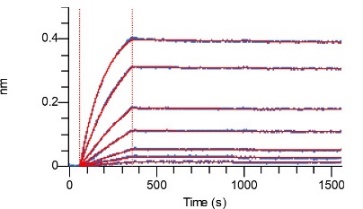 | 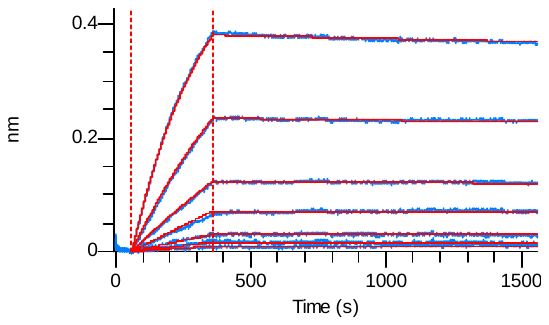 |

**Supplemental Figure 4A :** Binding affinity of VHH72-Fc improved clones to various RBD domains from different SARS-CoV variants: Bio-Layer Interferometry analysis of VHH-Fc immobilized proteins on anti-human Fc biosensors. Apparent binding kinetics of interaction between the VHH-Fc and the various RBD domains from SARS-CoV variants were evaluated in real time. Binding curves were fitted using a global 1:1 model.

|  | **RBD** | **RBD** |
| --- | --- | --- |
| **Mutants** | **SARS-CoV 2** | **SARS-CoV 1** |
| S57A | 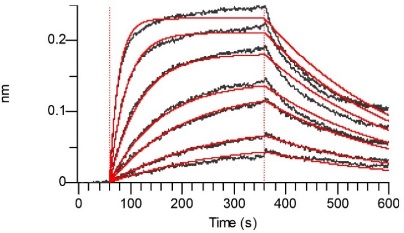 | 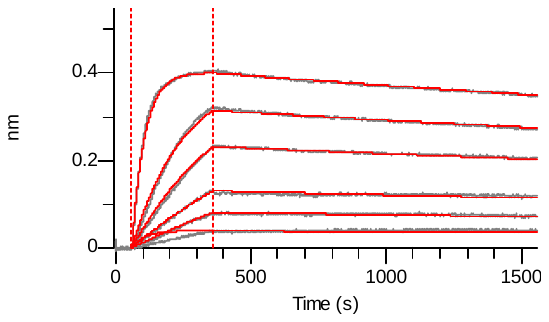 |
| S57G | 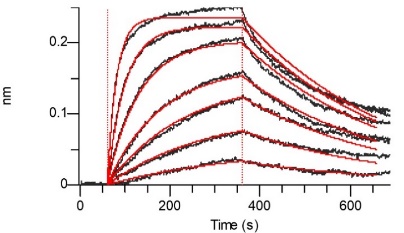 | 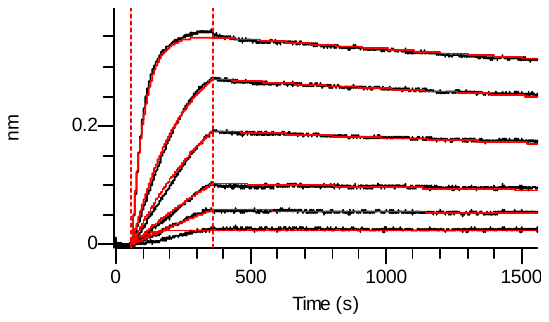 |
| T103V | 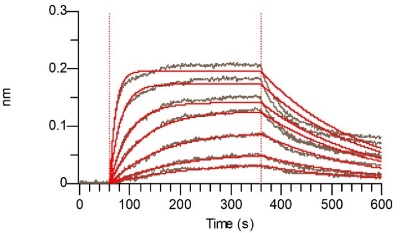 | 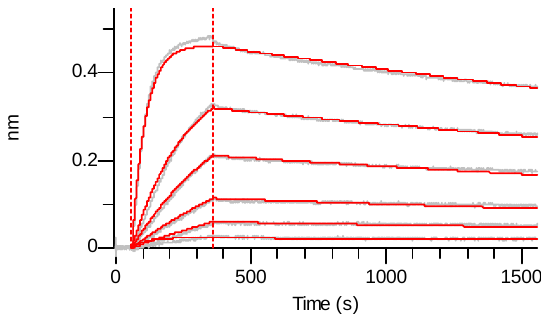 |
| V104W | 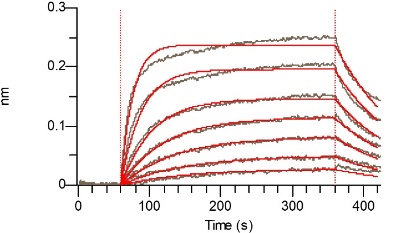 | 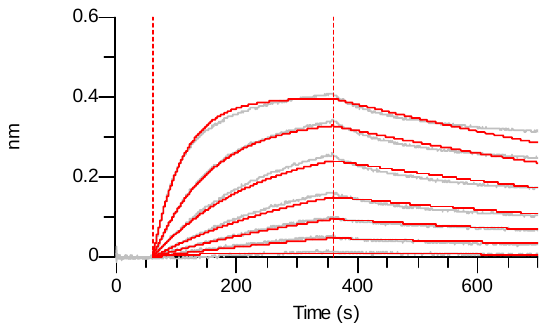 |
| S57G / T103V | 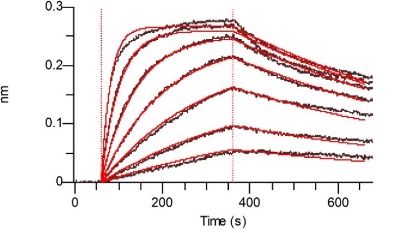 | 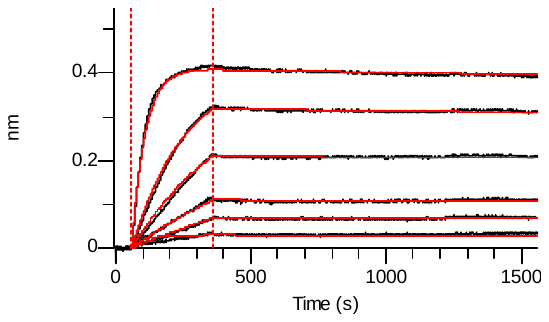 |
| S57G / V104W | 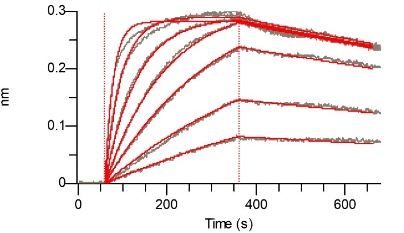 | 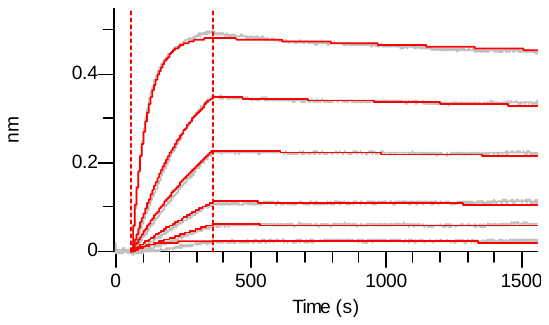 |
| T103V / V104W | 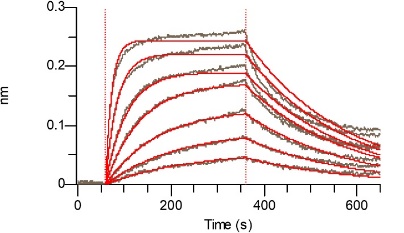 | 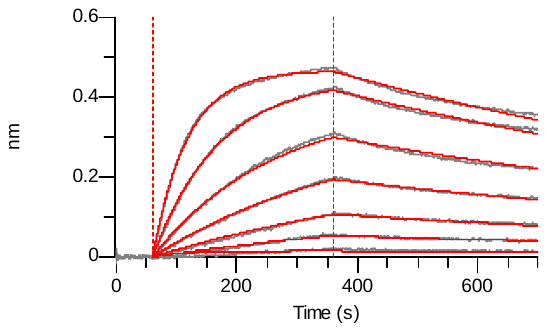 |
| S57G / T103V /V104W | 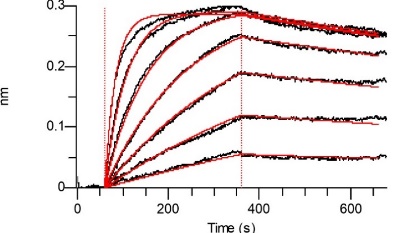 | 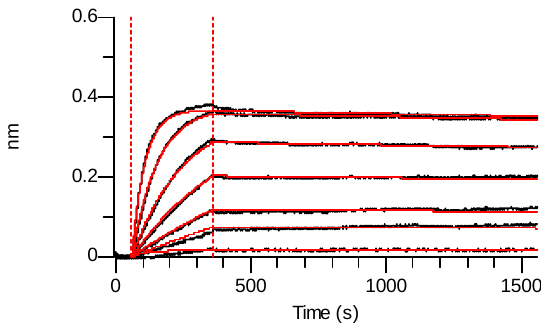 |
| VHH72 (WT) | 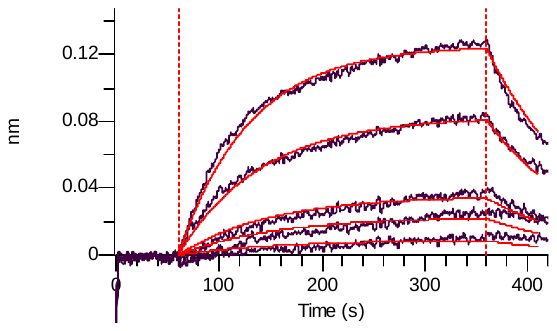 | 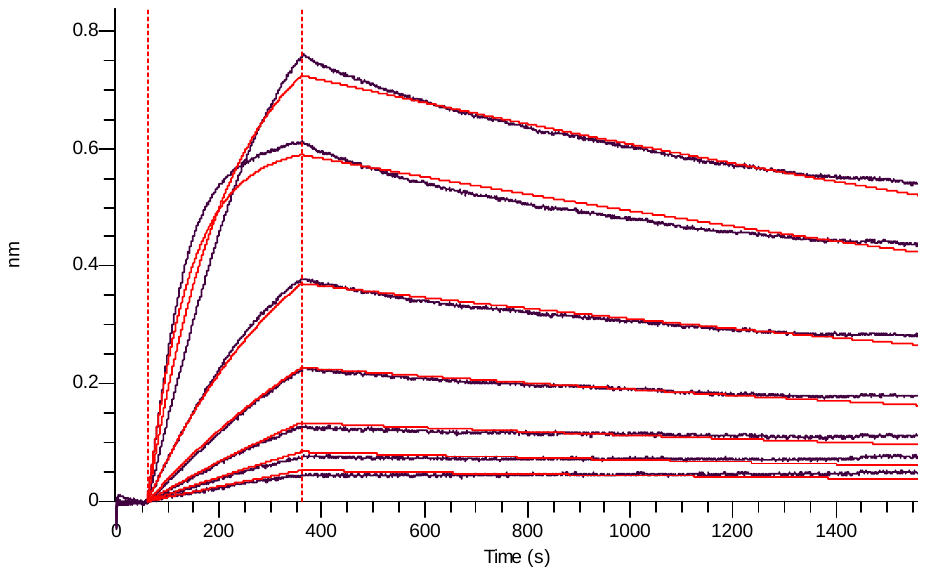 |

**Supplementary Figure 4B**

**Supplemental Figure 4B:** Binding affinity of VHH72-Fc monomutated clones to SARS-CoV 1 or SARS-CoV 2 variants: Bio-Layer Interferometry analysis of VHH-Fc immobilized proteins on anti-human Fc biosensors. Apparent binding kinetics of interaction between the VHH-Fc and the various RBD domains from SARS-CoV variants were evaluated in real time. Binding curves were fitted using a global 1:1 model.

|  | **RBD SARS-CoV 2** | | |
| --- | --- | --- | --- |
| **Mutation** | **K_D_ app (pM)** | **k_on_ (1/Ms)** | **k_off_ (1/s)** |
| S57A | 3460 | 1,07E+06 | 3,70E-03 |
| S57G | 3430 | 8,89E+05 | 3,05E-03 |
| T103V | 2970 | 1,64E+06 | 4,87E-03 |
| V104W | 2200 | 1,79E+06 | 3,94E-03 |
| S57G / T103V | 1550 | 9,21E+05 | 1,43E-03 |
| S57G / V104W | 528 | 1,08E+06 | 5,72E-04 |
| T103V / V104W | 3220 | 1,42E+06 | 4,58E-03 |
| S57G / T103V /V104W | 526 | 8,31E+05 | 4,37E-04 |
| VHH72 (WT) | 17460 | 5,73E+05 | 1,00E-02 |

**Supplementary Figure 4C**

|  | **RBD SARS-CoV 1** | | |
| --- | --- | --- | --- |
| **Mutation** | **K_D_ app (pM)** | **k_on_ (1/Ms)** | **k_off_ (1/s)** |
| S57A | 271 | 4,21E+05 | 1,14E-04 |
| S57G | 213 | 4,32E+05 | 9,18E-05 |
| T103V | 489 | 3,99E+05 | 1,95E-04 |
| V104W | 2620 | 3,72E+05 | 9,73E-04 |
| S57G / T103V | 54,23 | 4,14E+05 | 2,24E-05 |
| S57G / V104W | 134 | 3,53E+05 | 4,74E-05 |
| T103V / V104W | 2709 | 3,30E+05 | 8,95E-04 |
| S57G / T103V /V104W | 80 | 4,22E+05 | 3,35E-05 |
| VHH72 (WT) | 1143 | 2,41E+05 | 2,75E-04 |

**Supplemental Figure 4C:** Kinetics calculated values (K_D_ app; k_on and_ k_off_) of VHH72-Fc monomutated clones on SARS-CoV 2 and SARS-CoV 1.

**Supplementary Figure 5**

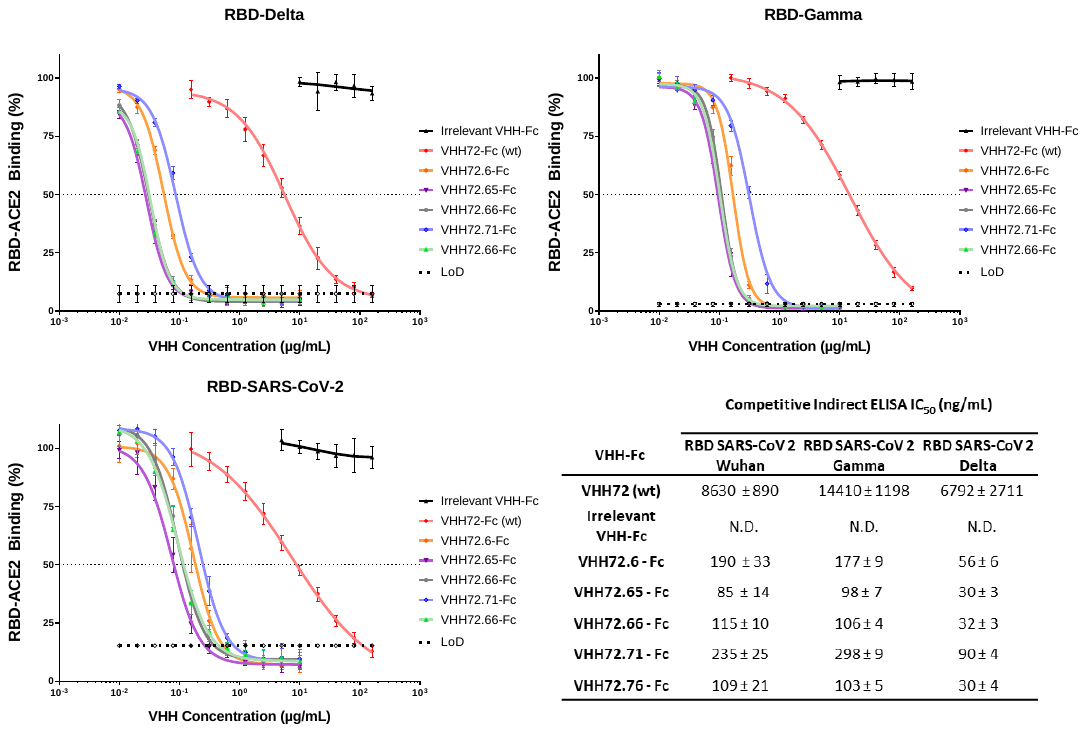

**Supplementary Figure 5:** Assessment of the ability of the selected VHH-Fc antibodies to block the interaction between ACE2 and the RBD domain of SARS-CoV-2 strain Delta, Gamma and Original in a competitive ELISA setup. Values represented correspond to three independent experiments.

**Supplementary Figure 6**

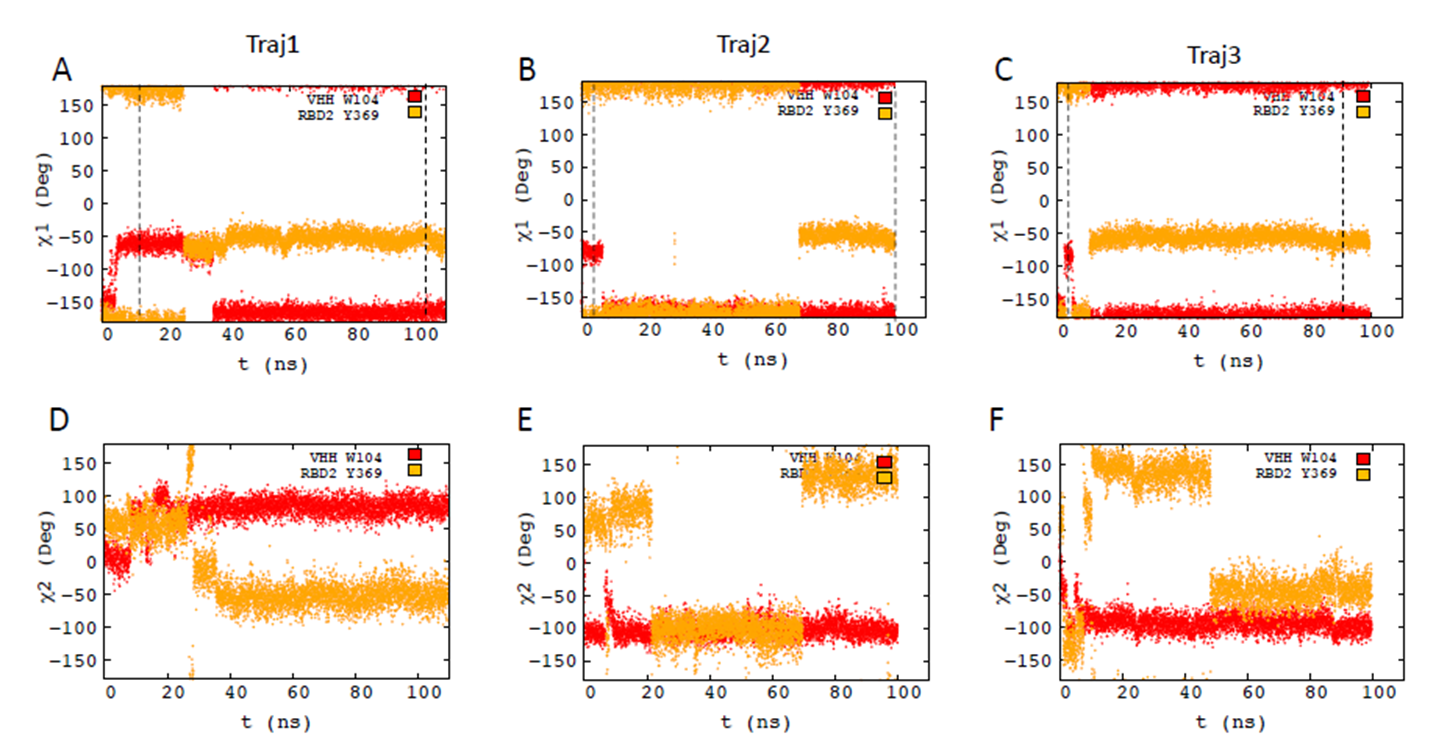

**Supplementary Figure 6: Orientation of SARS-CoV-2 RBD Y369 and VHH72 W104 along the MD trajectories**.

The χ1 dihedral angles of SARS-CoV-2 RBD Y369 (in red) and VHH72 W104 (in orange) are shown in A, B and C for trajectory 1, 2, 3. The χ2 dihedral angles of SARS-CoV-2 RBD Y369 (in red) and VHH72 W104 (in orange) are shown in D, E and F for trajectory 1, 2, 3.

**Supplementary Figure 7**

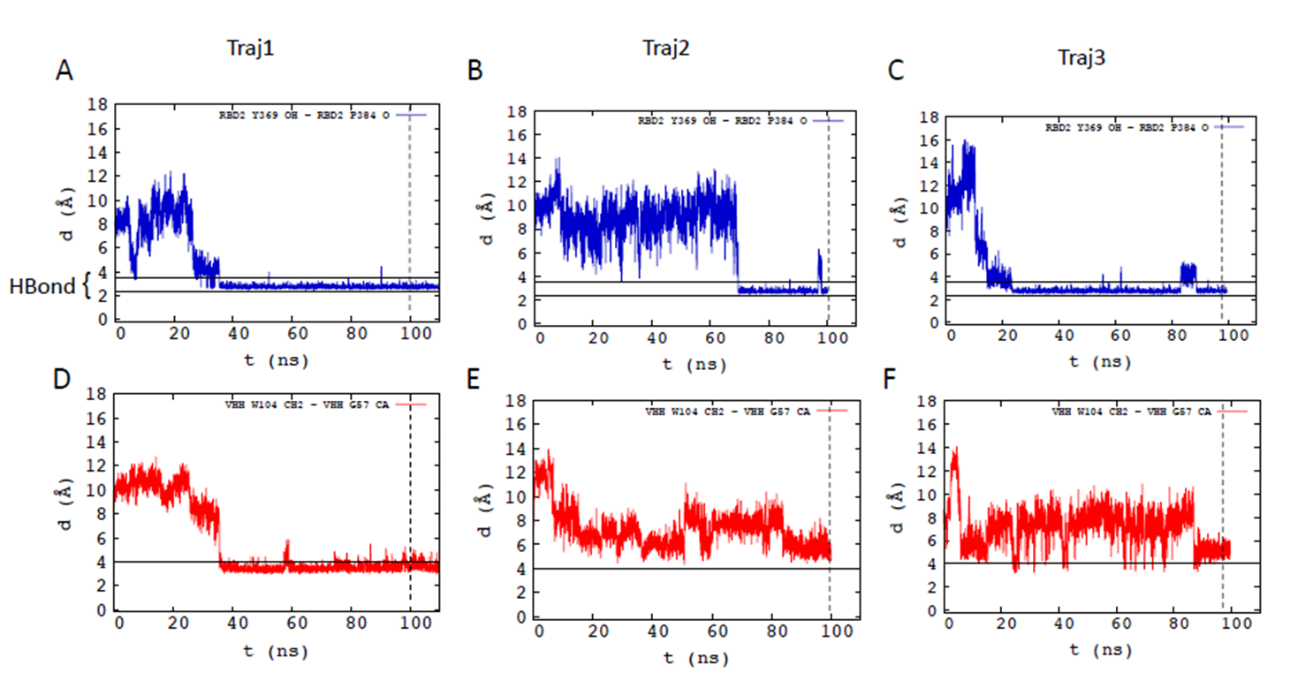

**Supplementary Figure 7: Interatom distances monitored along the three SARS Cov2 RBD Y369 and VHH72 W104 MD trajectories**.

The SARS Cov2 RBD Y369 OH-SARS-CoV-2 RBD P384 O distance is shown in A, B and C for trajectory 1, 2 and 3. The distances corresponding to a hydrogen bond are shown using black lines at y=2.3 Å and y=3.5 Å. The distance between the indole ring of VHH72 W104 and the Ca of residue 57 is shown in D, E and F for trajectory 1, 2 and 3.
